# Investigation of host plant contact with *Diaphorina citri* (Hemipreta: Psyllidae) by detecting *D. citri*-derived environmental DNA

**DOI:** 10.1101/2024.07.12.603183

**Authors:** T. Fujikawa, K. Fujiwara, H. Inoue, H. Hatomi, S. Hayashikawa

## Abstract

Citrus greening (Huanglongbing) disease has serious impacts on citrus production. Field monitoring of the Asian citrus psyllid *Diaphorina citri* (Hemiptera: Psyllidae), a vector of citrus greening disease, is essential to prevent the invasion and spread of this disease. This study reported a new method for determining the presence of *D. citri* and traces of contact with host plants by collecting leaves of host plants from the survey area and detecting the environmental DNA (eDNA) derived from *D. citri*. The results showed that the method can determine the presence of *D. citri.* As *D. citri-*derived eDNA is contained in DNA solutions of plants with a history of contact with *D. citri*, we succeeded in detecting not only genes conserved in *D. citri* mitochondria but also genes of *D. citri* symbiont organisms consisting of *Wolbachia*, *Wolbachia* phage, *Candidatus* Carsonella sp., and *Candidatus* Profftella sp. *Diaphorina citri*-derived eDNA could be detected in host plants even after only 10 min of contact with *D. citri* and could still be detected 6 months after contact. This technology has the potential to trace *D. citri* from their traces without individual detection and is expected to greatly contribute to the early detection and invasion warning of citrus greening disease in the future.

## Introduction

The Asian citrus psyllid *Diaphorina citri* Kuwayama (Hemiptera: Psyllidae) is a tiny phloem sap-sucking insect found in many parts of tropical and subtropical zones, mainly in Asia and the New World. Its hosts are mainly *Citrus* spp. and *Murraya* spp. (Sapindales: Rutaceae) and the vigorous multiplication of this insect can be observed on trees during the period when new leaves unfold. This insect is the principal vector of the citrus greening (Huanglongbing) disease, *Candidatus* Liberibacter asiaticus (CLas). This disease poses a serious threat to commercial citrus production worldwide (Grafton-Cardwell et al., 2013). *Diaphorina citri* acquires the pathogen by sucking the sap of an infected citrus tree and transmits the pathogen when it moves to a healthy citrus tree and sucks its sap. This transmission event is the primary step leading to the spread of citrus greening disease and is more likely to occur in the field than artificial grafting. Symptoms of citrus greening disease include blotchy chlorosis and/or mottling of leaves, yellowish shoots, vein corking, stunted growth, poor root growth, abnormal (small, green, and malformed) fruits, and death (Bové, 2006). As methods to treat or prevent this disease have not yet been established in the field, it is essential to control the insect vector *D. citri* in addition to removing pathogen-infected trees and preparing healthy nursery stocks (Grafton-Cardwell et al., 2013; Iwanami, 2022). Various insecticides with foliar or soil applications have been put into practical use for the control of *D. citri* and some have been successful in having strong insecticidal effects and reducing the proliferation of *D. citri* (Grafton-Cardwell et al., 2013; Iwanami, 2022). The effectiveness of insecticide treatment was confirmed by monitoring the presence of *D. citri* in the area by sampling *D. citri* using yellow sticky traps (Hall et al., 2010) and other methods, such as beating (Monzo et al., 2015). However, the bottleneck is whether *D. citri* can be visually identified as, when sampled, *D. citri* is often accompanied by many deteriorated insect bodies of other species. In our previous study, we developed a method to accurately confirm the presence of *D. citri* by extracting bulk DNA from many insects sampled using yellow sticky traps and performing polymerase chain reaction (PCR) targeting the mitochondrial cytochrome C oxidase subunit I (COI) gene specific to *D. citri* (Fujiwara et al., 2017). This method showed excellent performance for monitoring *D. citri* using yellow sticky traps. However, monitoring is only possible if sticky yellow traps are installed in advance in the field. In other words, if there is a new area where investigators would like to know about the presence of *D. citri*, they will need to set up yellow sticky traps in the area, collect the traps after leaving the traps for several weeks to several months, and then confirm the presence of *D. citri* from the bulk DNA extracted from the traps by PCR. Therefore, early and flexible monitoring is not possible. While setting the traps and leaving them for a certain period, *D. citri* can proliferate, and in some cases, provide the pathogen-acquired vector time to transmit the pathogen to healthy trees.

Environmental DNA (eDNA) refers to DNA derived from organisms present in the environment. By collecting and analyzing eDNA, we can assess the presence of specific species and the biodiversity of entire ecosystems. eDNA has been collected from bulk environmental samples across various ecosystems, such as marine, freshwater, sediment/soil, and air (Pawlowski et al., 2020; Roger et al., 2021; Johnson et al., 2022). This method has enabled the detection of species without directly sampling organisms, including invasive insects (e.g., Argentine ants in Japan) and alien fish (e.g., largemouth bass in China), thus allowing for the discovery of species in previously unexplored sites (Yasashimoto et al., 2021; Cheng et al., 2023). Traditional detection methods for *D. citri* extract DNA from individual insect samples. However, it is possible that DNA from the excretions of *D. citri* is present in the environment and can be extracted as eDNA. This offers the possibility of detecting this critical agricultural pest, which harbors the CLas-causing citrus greening disease, and raises an intriguing question about the technical feasibility of using molecular tools to detect the presence of even small insects through eDNA.

In this study, we report a new method for determining the presence of *D. citri* and traces of contact with host plants by collecting leaves of host plants from the survey area and detecting the eDNA derived from *D. citri*.

## Materials and Methods

### Collection and DNA extraction of *D. citi* and plant samples

Ten individuals of *D. citri* used for genome sequencing were collected from Ogimi Village, Okinawa Prefecture, Japan, and fixed with absolute ethanol. DNA was extracted from these samples using the DNeasy Blood and Tissue kit (Qiagen, Hilden, Germany). *Diaphorina citri* used for the PCR assay were collected from various locations in Japan: Nago City and Ogimi Village of Okinawa Island; China Town of Okinoerabujima Island; Amami City of Amami-Oshima Island; Yoron Town of Yoron Island; and Isen Town of Tokunoshima Island. Various sternorrhynchous Hemiptera (Psylloidea, Aphidoidea, and Coccoidea) were collected from the warm southwestern area of the Satsuma Peninsula and the Amami Islands in Kagoshima Prefecture, Japan (Suppl. Tables 1 and 2). Leaves of *Murraya paniculata* plants with a history of *D. citri* infestation were collected in the experimental greenhouse of the Oshima Branch, Kagoshima Prefectural Institute for Agricultural Development on Amami-Oshima Island, Kagoshima Prefecture, Japan. Also, Healthy (*D. citri*-free) leaves of *M*. *paniculata* were collected in a *D. citri*-free experimental greenhouse of the Institute for Plant Protection, NARO, Tsukuba City, Ibaraki Prefecture, Japan (*D. citri* do not live in this area). Leaves of *M*. *paniculata* and *Citrus* spp. in the field were arbitrarily selected from the trees where *D. citri* was collected or from the surrounding trees that were not clearly (visually) infested with *D. citri*. All samples were fixed with ethanol to kill pests and pathogens. The collected leaves were wiped with paper soaked in 70% ethanol to remove dirt. DNA was extracted from *D. citri* (except for genome sequencing), insects, and plant leaves using the Macherey-Nagel NucleoSpin Food Kit (Takara Bio Inc., Shiga, Japan).

### Draft genome sequencing of *D. citri* and its symbionts

To obtain the partial genome sequences of *D. citri* and its symbionts, genomic DNA was sequenced from ten individuals of *D. citri* collected from citrus trees in Ogimi Village, Okinawa Prefecture, Japan. Genomic DNA was extracted from the body and fixed in ethanol. DNA libraries were prepared from genomic DNA using an Ion Plus fragment library kit, with physical shearing and size selection (approximately 200 bp), and were sequenced using an Ion PGM sequencer with the Ion PGM Hi-Q View OT2 kit, Ion PGM Hi-Q View sequencing kit, and Ion 318 Chip Kit v2 (all from Thermo Fisher Scientific, Inc., Waltham, MA, USA), according to the manufacturer’s instructions. The sequence reads were evaluated for quality (quality scores ≥20), and adapter sequences were trimmed using the CLC Genomics Workbench v12 (Qiagen). The obtained reads were mapped to the reference sequence of the *D. citri* genome (GenBank Accession number: GCA_000475195.1). Reads that were not mapped to *D. citri* genome were collected and assembled *de novo* into contigs using the CLC Genomics workbench with default parameters (mapping mode = create simple contig sequences [fast], automatic bubble size = yes, minimum contig length = 500, automatic word size = yes, performing scaffolding = yes, auto-detect paired distances = yes). These contigs were subjected to a BLAST search, and symbiont organism sequences were selected. In addition, the representative genome sequences of symbiont candidates obtained from the BLAST search results were prepared as references, and the reads from the genomic DNA of *D. citri* including the symbionts, were mapped to the references.

### PCR primers and conditions

For the detection of *D. citri* and its symbiont organisms, respective primer sets were designed (Table 1). Specific primer sets for *D. citri* were designed from the sequences of the 12S rDNA, cytochrome C oxidase subunit I (COI), and ND4 (NADH-ubiquinone oxidoreductase chain 4) genes, with reference to the mitochondrial genome (GenBank Accession number MG489916). Specific primer sets for symbionts were designed with reference to the corresponding contigs obtained from the DNA of ten *D. citri* individuals (Suppl. Text 1). Contig_136 for *Wolbachia* spp., Contig_148 for *Wolbachia* phages, and Contig_207 for *Ca.* Carsonella spp., Contig_27 for *Ca.* Profftella spp. was also used in this study. These primer sets were designed to amplify products of approximately 150–250 bp so that they could be used not only for conventional (end-point) PCR (cPCR) but also for quantitative PCR (qPCR) using an intercalator (SYBR green). PCR was performed as described in our previous report (Fujiwara et al., 2017) with some modifications. Namely, cPCR was performed in a 20-μl reaction mixture with 1× PCR buffer containing 2 μl of extracted DNA, 0.2 mM deoxyribonucleotide triphosphates, 0.5 U *ExTaq* HS polymerase (Takara Bio Inc.), and 0.25 μM of the respective primer sets listed in Table 1. The PCR reaction mixtures were heated at 94LJ for 4 min, followed by 40 cycles of denaturation at 94LJ for 30 s, annealing at 60LJ for 30 s, extension at 72LJ for 30 s, and a final extension at 72LJ for 7 min. Ten microliters of PCR products with 2 μl of RE-DYE (Toyobo, Osaka, Japan) was analyzed using 1.5% agarose gel electrophoresis containing 0.5 mg/ml ethidium bromide. The qPCR was performed in a QuantStudio 3 real-time PCR system (Thermo Fisher Scientific Inc.) using a TB green *Premix ExTaq* II (Tli RNaseH Plus) kit (Takara Bio Inc.) to label and amplify templates in accordance with the manufacturer’s protocol. The qPCR reaction mixtures were heated at 94LJ for 4 min, followed by 42 cycles of denaturation at 94LJ for 30 s, annealing at 60LJ for 30 s, extension at 72LJ for 30 s, and a final extension at 72LJ for 7 min. Then, for melt curve analysis, after denaturation of 94°C for 30 s, the mixtures were incubated at 60°C for 1 min and the melting temperature was monitored while increasing the temperature up to 94°C.

**Table 1.**
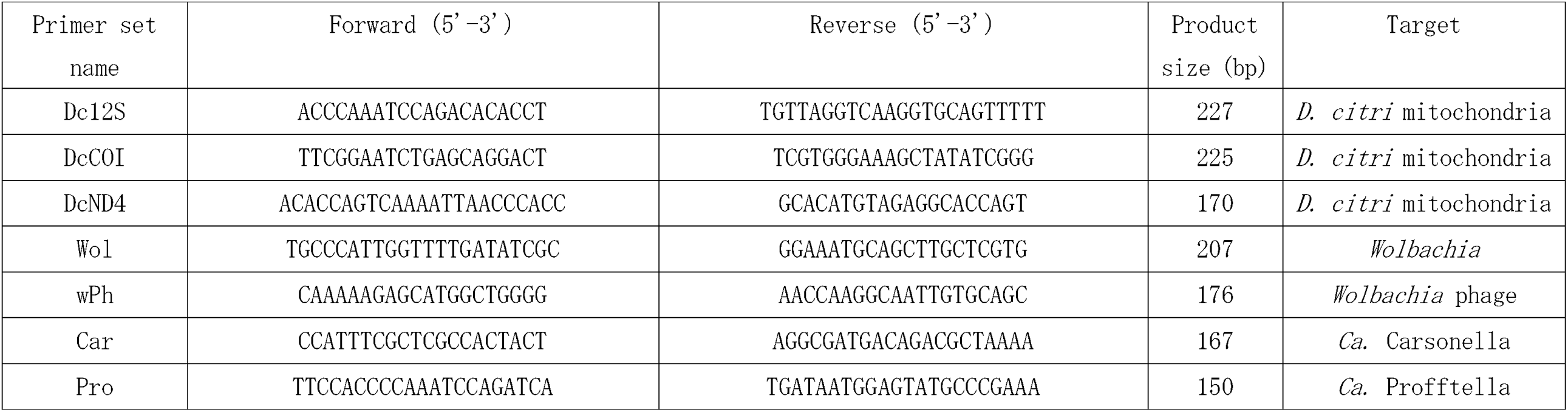
Primers for detecting D. citri and its symbionts used in this study

### *Diaphorina citri* contact duration test capable of detecting *D. citri-*derived eDNA

Thirty-two *M. paniculata* seedlings (10–15 cm in height) were grown from seeds for this test in an environment isolated from *D. citri*. In contrast, several *M. paniculata* trees were set up in a closed environment where *D. citri* was released and allowed to multiply sufficiently. For each seedling, ten individuals of adult *D. citri* were attached to the leaves and covered with plastic bags to prevent *D. citri* from escaping. Small air holes are created in the bags. *Diaphorina citri* inoculation times were set as follows: 10 min, 30 min, 1 h, 2 h, 3 h, 24 h, and 45 h, and bipinnately compounded leaves were collected at each time interval. Seedlings without *D. citri* (non-inoculated) were used as a control. After each compound leaf was collected, DNA was extracted. PCR was performed using *D. citri* primers listed in Table 1. The test was performed with four replicates.

### Residual duration test of *D. citri*-derived eDNA

Ten *Muraya* seedlings were used (each seedling had at least nine compound leaves). They were grown from seeds in an environment isolated from *D. citri*. In addition, *D. citri* was propagated and prepared in large quantities, as mentioned above. For each seedling, 100 adult *D. citri* were attached to the leaves and covered with a fine-mesh net bag as described above. *Diaphorina citri* inoculation times were set at 24 h and 1 week. As a control, compound leaves were collected from seedlings before inoculation with *D. citri* (before inoculation). After the inoculation period, *D. citri* was completely removed, and the seedlings were kept covered with bags for 0, 1, 3, 7, 14, 30, 60, 90, and 180 d. New shoots that developed after the end of *D. citri*-inoculation were removed. Compound leaves were collected at each time point and were extracted for qPCR using *D. citri* primers listed in Table 1. The test was performed with five replicates.

## Results

### Obtaining DNA sequences of *D. citri* and its symbionts

Draft genomic DNA of *D. citri* was sequenced from ten individuals using the next-generation sequencer Ion PGM. This genomic DNA may include symbiont organisms of *D. citri.* The number of short sequence reads obtained varied among individuals; however, the average length was similar (Suppl. Table 3). When each of these reads was mapped to the reference *D. citri* genome, reads ranging from 91.83% to 94.40% were fixed. The read-fixed location of the *D. citri* genome contains a mitochondrial genome containing 12S rDNA, cytochrome C oxidase subunit I (COI), and NADH-ubiquinone oxidoreductase chain 4 (ND4) genes. Sequence reads that did not map to the *D. citri* genome were obtained from all individuals. Thus, these reads were assembled and 62 contigs (average length = 984.61 bp) were obtained. When these contig sequences were submitted to the NCBI BLAST search, they matched the bacterial and phage sequences of some contigs (Suppl. Table 4). These sequences appear to be derived from *D. citri* symbionts. Thus, *Wolbachia* spp., *Candidatus* Carsonella spp., *Candidatus* Profftella spp., and *Wolbachia* phages are listed as *D. citri* symbiont candidates. In addition, when we prepared the reference genomes (GenBank accession numbers AM999887 and CP034335 for *Wolbachia* spp. and CP012411 for *Ca*.

Carsonella spp.; CP012591 for *Ca.* Profftella spp.; KX522565 for *Wolbachia* phage) that were closely related to each of the four symbiont candidates and mapped the reads of each *D. citri* individual, all of which were fixed to the four references (Suppl. Fig. 1). In contrast, the BLAST results of contigs hit organisms other than the four symbionts but the presence or absence of these DNA sequences varied among each individual of *D. citri*, we excluded them from the essential symbionts of *D. citri*. Therefore, we confirmed that the DNA of each *D. citri* individual contained DNA from four symbiont organisms.

### Detection of *D. citri* and its symbiont organisms using new primers

By sequencing the *D. citri* genome, we were able to obtain not only the *D. ctiri* genome but also the DNA sequences of the symbiont organisms; therefore, we designed primers that can specifically detect *D. citri* and its symbiont organisms. To evaluate the performance of these PCR primers, we first performed PCR using the DNA of *D. citri* collected from various parts of the Southwestern islands of Japan as templates, resulting in amplicons obtained from all *D. citri* used in this test (Fig. 1). Using these primers, we confirmed that the target genes were amplified from *D. citri* DNA, regardless of the region or plants from which they were collected and that the four symbiont organisms were universally present in *D. citri*. Next, we investigated whether the *D. citri* primers, in particular the *D. citri* symbiont primers, could be used regardless of the growth stage of *D. citri*, and found that the target genes were amplified from the DNA of both adults and nymphs. In addition, we performed PCR on the DNA of *Murraya* that had been infested with *D. citri* (Suppl. Fig. 2) and found that the results were positive not only for the mitochondrial genome of *D. citri* (Fig. 2A, B, and C) but also for the symbionts (Fig. 2E and G).

**Figure 1.**
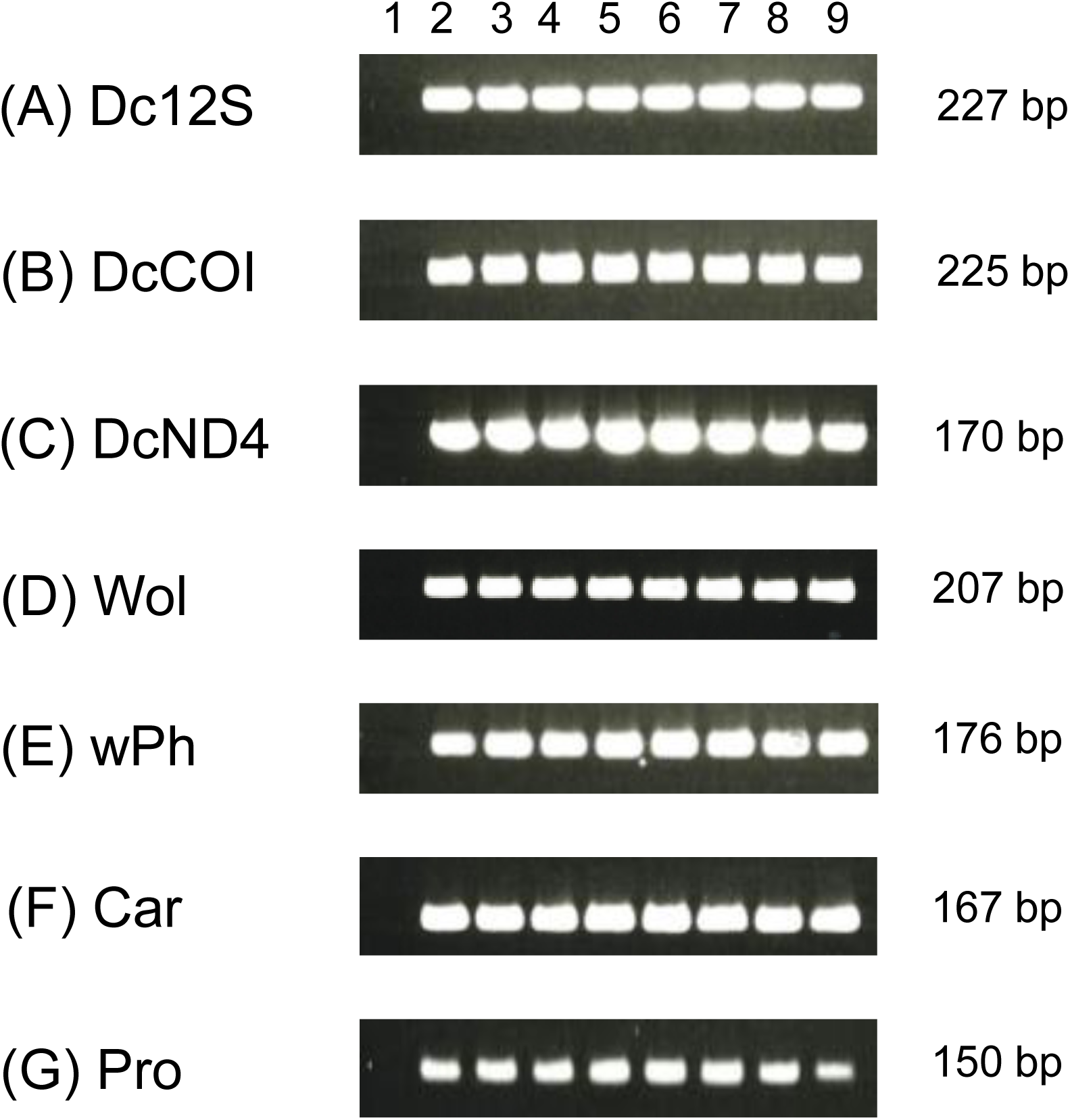
PCR results using each primer with the DNA of *D. citri* in Japan. When DNA was extracted from *D. citri* and used as a template for PCR using each primer, a single amplification product with the expected amplification size was obtained for all *D. citri* in this test. The targets of (A) Dc12s, (B) DcCOI, and (C) DcND4 are the mitochondrial genome of *D. citri*. The target of (D) Wol is *Wolbachia* spp., that of (E) wPh is *Wolbachia* phage, that of (F) Car is *Ca.* Carsonella spp., and that of (G) Pro is *Ca.* Profftella spp. Each template is as follows, Lane 1: water (negative control); Lanes 2 and 3: *D. citri* collected from *Murraya* trees in Nago city, Okinawa Prefecture; Lane 4: that from *Citrus depressa* tree in Nago city; Lane 5: that from *Murraya* trees in China town (Okinoerabujima island), Kagoshima Prefecture; Lane 6: that from *Murraya* trees of the experimental greenhouse of Oshima Branch Kagoshima Prefectural Institute for Agricultural Development in Amami city (Amami-Oshima island), Kagoshima Prefecture; Lane 7: that from *Murraya* trees in Yoron town (Yoron island), Kagoshima Prefecture; Lane 8: that from *Murraya* trees in Isen town (Tokunoshima island), Kagoshima Prefecture; and Lane 9: that from *citrus* trees in Ogimi village, Okinawa Prefecture (same as one of the sequenced samples).

**Figure 2.**
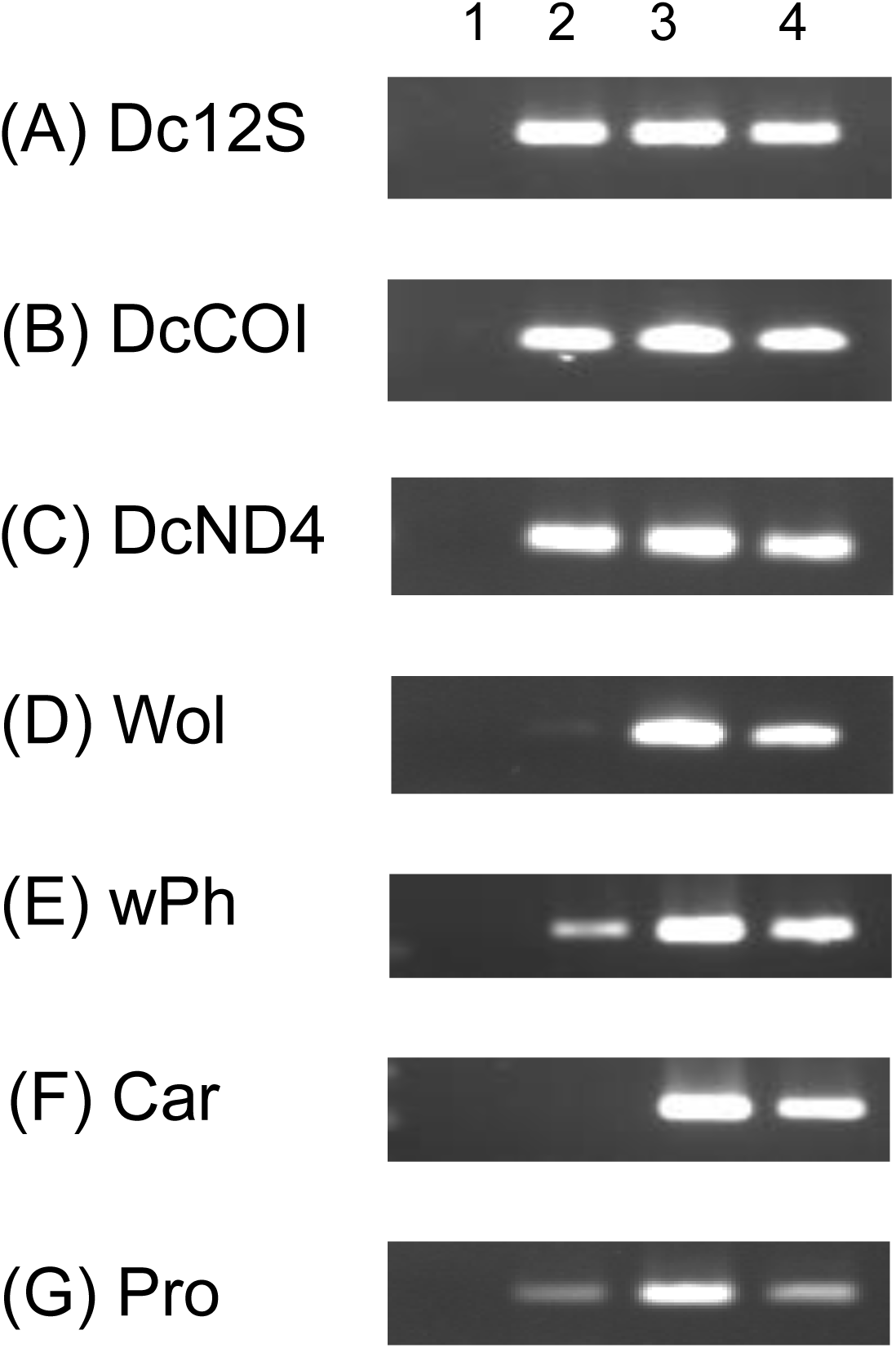
PCR results using each primer with the DNA of host leaves that *D. citri* had been parasitized, adults and nymphs of *D. citri*. From *Murraya* infested with *D. citri* like Suppl Fig 2, five adult individuals, five nymph individuals, and three leaves of *Murraya* tree (whose insect bodies had been removed and the surface had been washed with ethanol) were collected and DNA extracted. PCR was performed using each primer set with each template DNA. The targets of (A) Dc12s, (B) DcCOI, and (C)DcND4 are the mitochondrial genome of *D. citri*. The target of (D) Wol is *Wolbachia* spp., that of (E) wPh is *Wolbachia* phage, that of (F) Car is *Ca.* Carsonella spp., and that of (G) Pro is *Ca.* Profftella spp. Each template is as follows, Lane 1: water (negative control); Lane 2: leaves; Lane 3: adults of *D. citri*; and Lane 4: nymphs of *D. citri*.

In the present study, we investigated whether *D. citri* primers could identify plants with a history of *D. citri* infestation. *Diaphorina citri* was allowed to roam freely for approximately six months to a year in the experimental greenhouse of the Oshima Branch, Kagoshima Prefectural Institute for Agricultural Development (Amami-Oshima Island) where multiple *Murraya* trees were prepared. Amami Oshima, where the experimental greenhouse is located, is in a subtropical zone, and *D. citri* can grow and multiply throughout the year. During this period, we confirmed that *D. citri* grew sufficiently and repeatedly on *Murraya* trees. Next, we secured the trees from which *D. citri* had been removed, visually inspected the leaves that did not contain *D. citri*, extracted DNA, and used them as templates for PCR. As a control, *D. citri*-free *Murraya* trees were prepared in a greenhouse at the Institute for Plant Protection, NARO, Tsukuba City (an area that *D. citri* cannot inhabit), and DNA was extracted as a template for PCR. Examples of Leaves with or likely to have a history of *D. citri* infestation and *D. citri*-free leaves are shown in Suppl. Fig.3. *Diaphorina citri* was detected using seven primer sets in both adults and nymphs (except that DNA from nymphs was not PCR-amplified using DcCOI in this figure) and there was variation in the amplification of DNA from leaves in lanes 4 to 11 (Fig. 3). Thus, using multiple primers, we confirmed that *D. citri*-derived eDNA were amplified, and as a result, we were able to understand the infestation history of *D. citri*. PCR amplification did not occur with any of the primers in *D. citri*-free leaves in lane 12 (Fig. 3). Similar results were confirmed using qPCR (Table 2). Amplification products were obtained using all seven primers with DNA from adult *D. citri* and the Ct values were determined. Target amplification was evaluated using the Tm value and even if a Ct value was obtained, if the Tm value was different, it was considered non-detection. qPCR using *D. citri* primers yielded specific amplification products in *Murraya* leaves with a history of *D. citri* infestation (although Wol was not detected in some cases). Furthermore, when the *D. citri* primer sets designed in this study are used to detect the trace of *D. citri* in plants in the field, there is a possibility that related species other than *D. citri* may be erroneously detected. Therefore, among the small hemipterans collected in areas in Japan where *D. citri* are present or may be found because of the expansion of their distribution, we performed qPCR or cPCR using *D. citri* primers with the DNA as templates for various Psylloidea species, which cannot multiply on Rutaceae plants but may temporarily visit them, and other small Hemiptera species, which feed on *Citrus* and *Murraya* trees. qPCR using *D. citri* primers (DcND4, Wol, and wPh) sometimes yielded false-positive results in the Psylloidea group (Suppl. Table 1). In the cPCR assay for small Hemiptera collected from *Citrus* and *Murraya* trees from the fields of the warm southwestern regions in Japan, *D. citri* primers targeting the mitochondrial genome could detect *D. citri* (However, CIC01 and CIC03 of Cicadellidae were also detected) and *D. citri* symbiont primers targeting *Wolbachia* phage, *Ca*. Carsonella, and *Ca*. Profftella detected *D. citri* with high specificity (Suppl. Fig. 4).

**Figure 3.**
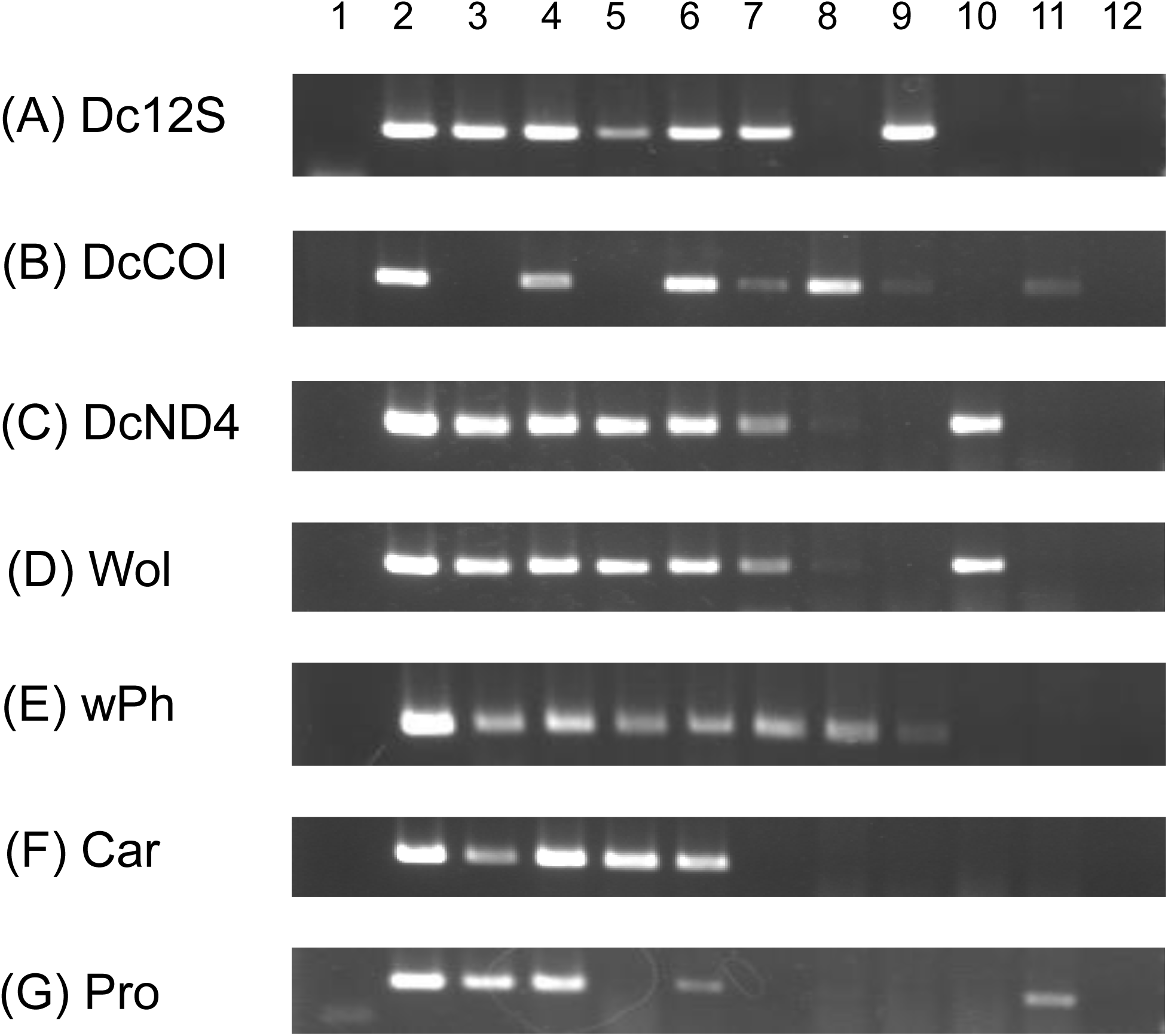
PCR results using each primer with the DNA of *Murraya* leaves with a history of *D. citri* infestation. Leaves of *Murraya* spp. with a history of *D. citri* infestation in the experimental greenhouse were sampled for detecting *D. citri*-derived environmental DNA. The targets of (A) Dc12s, (B) DcCOI, and (C) DcND4 are the mitochondrial genome of *D. citri*. The target of (D) Wol is *Wolbachia* spp., that of (E) wPh is *Wolbachia* phage, that of (F) Car is *Ca.* Carsonella spp., and that of (G) Pro is *Ca.* Profftella spp. Each template is as follows, Lane 1: water (negative control); Lane 2: an individual adult of *D. citri*; Lane 3: an individual nymph of *D. citri*; Lane 4 to 11: a group of three leaves of *Murraya* that have the history of *D. citri* infestation; Lane 12: a group of three leaves of *D. citri-*free *Murraya*.

**Table 2.** Ct value of quantitative PCR for detecting *D. citri-*derived environmental DNAs

|  | *D. citri*<br>adult | *Murraya*-Dc<br>-1 | *Murraya*-Dc<br>-2 | *Murraya*-Dc<br>-3 | *Murraya*-health<br>y-1 | *Murraya*-health<br>y-2 | *Murraya*-health<br>y-3 | None |
| --- | --- | --- | --- | --- | --- | --- | --- | --- |
| Dc12S | 21.40 | 32.81 | 31.55 | 32.59 | Diff<sup>1</sup> | Diff | Diff | ND<sup>2</sup> |
| DcCOI | 21.01 | 30.94 | 31.39 | 33.42 | Diff | ND | ND | ND |
| DcND4 | 20.86 | 31.79 | 31.01 | 32.32 | ND | ND | ND | ND |
| Wol | 23.92 | 37.01 | ND | ND | Diff | ND | ND | ND |
| wPh | 24.93 | 33.97 | 34.38 | 35.63 | ND | ND | ND | ND |
| Car | 26.54 | 34.85 | 32.96 | 36.94 | ND | ND | ND | ND |
| Pro | 23.71 | 32.89 | 31.43 | 34.87 | ND | ND | ND | ND |

*D. citri* adult: an individual adult of *D. citri*; *Murraya*-Dc-1 to Dc-3: a group of three leaves of *Murraya* that have the history of *D. citri* infestation; *Murraya*-healthy-1 to Healthy-3: a group of three leaves of *D. citri*-free *Murraya*.
The T<sub>m</sub> values (°C) of each PCR product are as follows: Dc12s, 75.99±0.09; DcCOI, 77.41±0.30; DcND4, 76.55±0.37; Wol, 78.75±1.27; wPh, 79.91±0.33; Car, 80.99±0.30; Pro, 75.10±0.36.
<sup>1</sup>Diffrent T<sub>m</sub> value: Since the T<sub>m</sub> value is different from the expected T<sub>m</sub> value of amplification product, it is determined that it is not detected.
<sup>2</sup>Not detected: No amplification product was obtained.

### *Diaphorina citri* contact duration test

*Diaphorina citri*-derived eDNA was detected in plants with a history of *D. citri* infestation. Therefore, we investigated the duration for which *D. citri* must be in contact with host leaves before *D. citri*-derived eDNA can be detected. Ten *D. citri* were attached to the leaves of *Murraya* seedlings, covered with bags, and left for a certain period for *D. citri* inoculation (Suppl. Fig. 5). Thereafter, the leaves were collected over time, DNA was extracted, and cPCR was performed using *D. citri* primers (Suppl. Fig. 6). The cPCR results confirmed that the DNA contained *D. citri*-derived eDNA (Suppl. Fig. 6). The detection results (presence or absence of amplified products) were recorded at each inoculation time (Fig. 4). No *D. citri*-derived eDNA was detected in seedlings to which *D. citri* was not attached (not-inoculated). Even after an inoculation time of 10 min, trace DNA of *D. citri* could be detected. However, as the inoculation time increased, the number of genes detected also increased, reaching its highest number at 2 h after inoculation. Furthermore, even if PCR using all seven primer sets did not simultaneously yield positive results at the same time, a history of contact with *D. citri* was suspected if one of the primers was positive.

**Figure 4.**
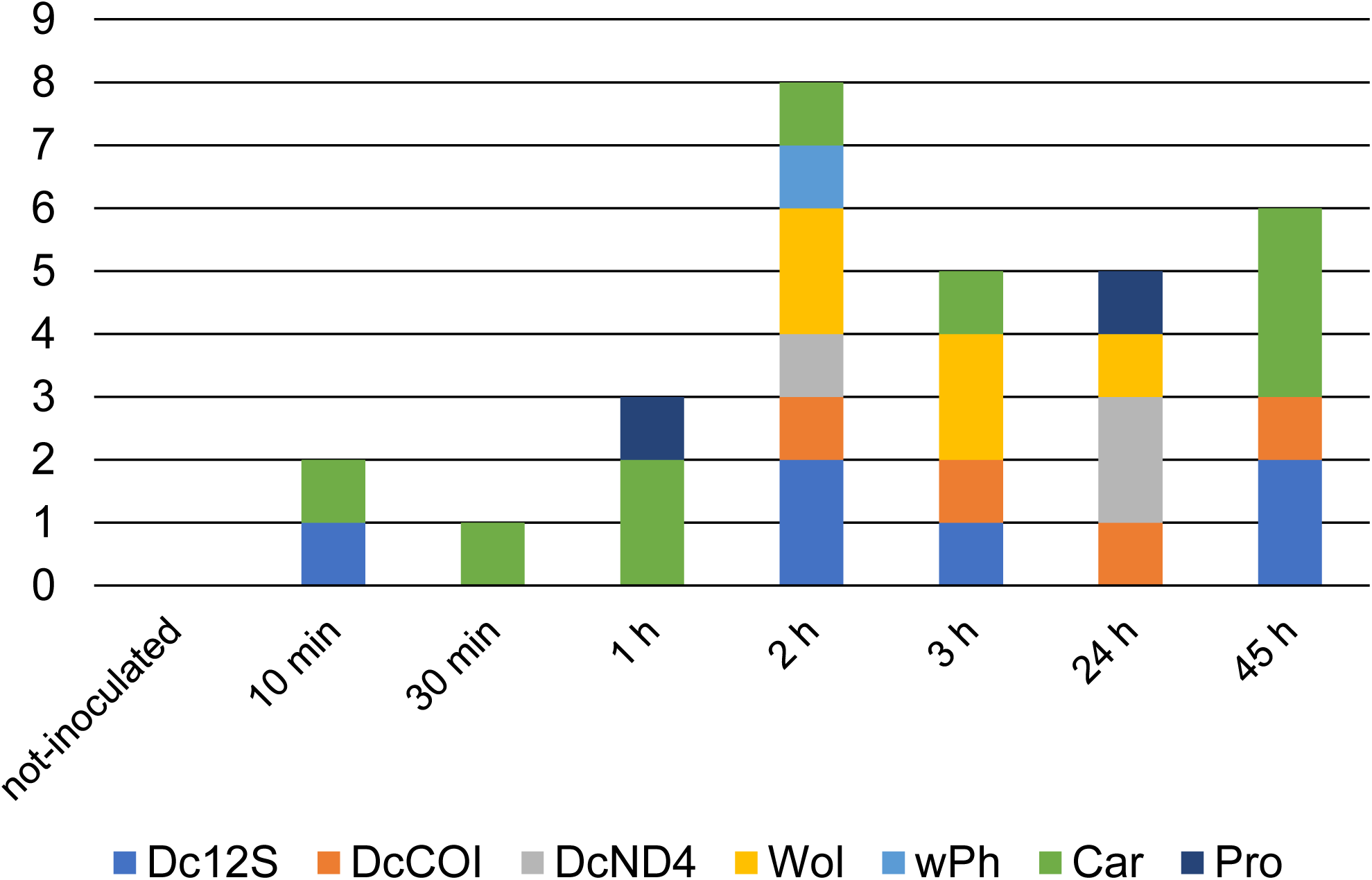
Total number of *D. citri*-derived environmental DNA detected at different ***D. citri* inoculation times.** DNA from *Murraya* leaves, which were inoculated with *D. citri*, were extracted and used for cPCR using *D. citri* primers. The presence or absence of PCR amplification products using each primer (Suppl. Fig. 6) was summed for each different inoculation time. In this figure, the number of detections for each environmental DNA at each inoculation time was accumulated. Number of trees tested (n)= 4.

### Residual duration test

We found that if *D. citri* attached to *Murraya* for a short period, it left traces on its leaves as *D. citri*-derived eDNA. Next, we investigated how long the eDNA remained. Each *Murraya* seedling was infested with 100 individuals of *D. citri*, covered with a fine-mesh net bag. After the inoculation period (1 or 7 d), *D. citri* was completely removed and the leaves inoculated with *D. citri* were kept covered with bags for a certain period (Suppl. Fig. 7). The leaves were collected at each time point, DNA was extracted, and qPCR was performed using *D. citri* primers (Suppl. Table 5; Suppl. Table 6). When the Ct values were obtained by qPCR, it was considered “detected” and the average of the total number of “detected” was expressed for each residual duration time (Fig. 5). There was no significant difference between the inoculation times of 1 and 7 d, and the number of detections decreased as the residual duration increased. The study examined traces for up to 180 days (six months), and some *D. citri*-derived eDNA were still detectable during that period.

**Figure 5.**
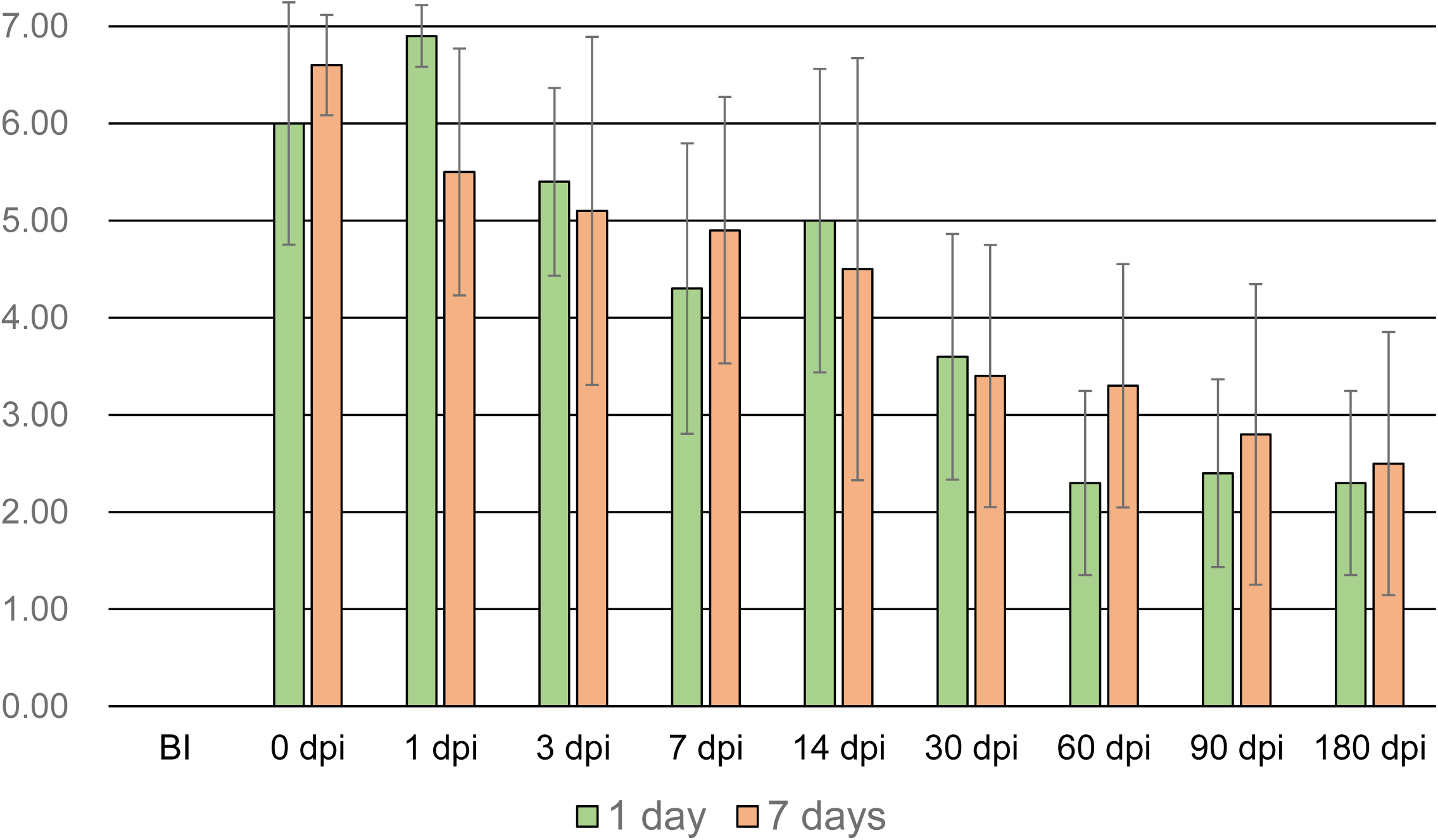
Total number of *D. citri-*derived environmental DNA detected at different residual durations. *Muraya* leaves inoculated with *D. citri* for 1 or 7 days were maintained from 0 to 180 days (0, 1, 3, 7, 14, 30, 60, 90, and 180 days), after which DNA was extracted and used for qPCR using *D. citri* primers. When the Ct values (Suppl. Table 5) were obtained for the amplified products by qPCR using each primer, it was considered a “detected” and the detected number was summed for each residual duration (Suppl. Table 6). In this figure, the average values of the total number of “detected” for each residual duration after *D. citri* was inoculated for 1 day or 7 days are shown. The vertical axis is the total number of “detected”. The maximum value is 7. In each sample, n=10. Error bars indicated standard deviation. BI: before inoculation. There is no significant difference between the inoculation times of 1 and 7 days.

### Adaptability in *D. citri* habitat areas

Because *D. citri*-derived eDNA could be detected in plant leaves in the greenhouse, and it was possible to detect *D. citri* sufficiently, we investigated its adaptability in the field. We collected *Citrus* and *Murraya* leaves from various locations on Okinoerabujima Island, where *D. citri* inhabited and citrus greening disease occurs, to investigate whether *D. citri*-derived eDNA from these leaves could be detected. When cPCR was performed on the 24 samples, one or more types of *D. citri*-derived eDNA were detected in multiple samples (Table 3; Suppl. Fig. 8). Two samples in which *D. citri* were found near the leaves (on the same tree or adjacent trees) were also included among the samples in which eDNA was detected (Table 3). Similarly, qPCR was performed on 35 samples, which showed Ct values for *D. citri* and its endophytic bacterial genes (Table 4). Particularly, PCR using DcCOI or DcND4 yielded positive results in many samples. In addition, for sample “1-2 of *Citrus* location J,” the PCR results were positive with all primer sets. These results confirmed that *D. citri*-derived eDNA was detected even in the on-site leaves.

**Table 3.**
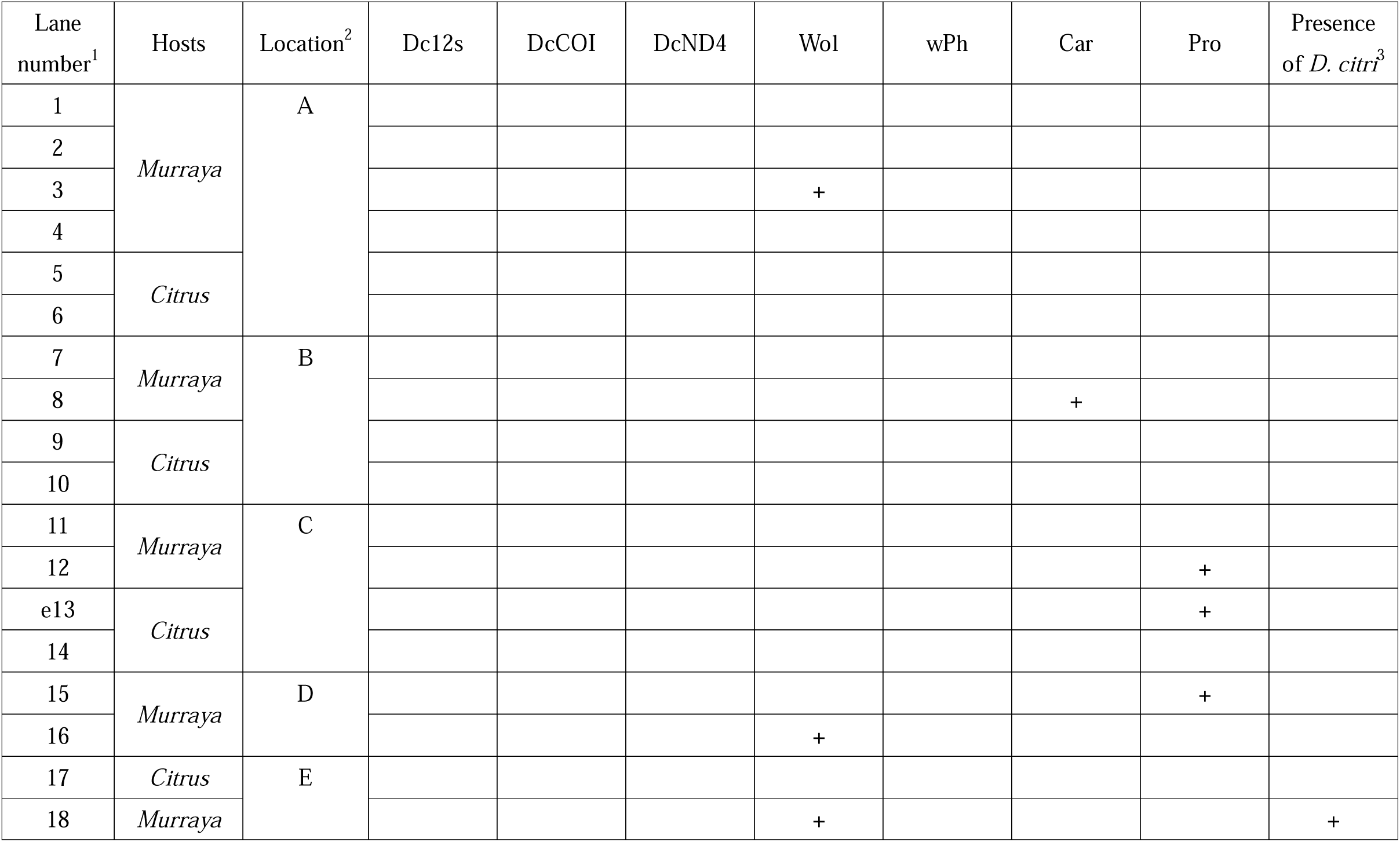
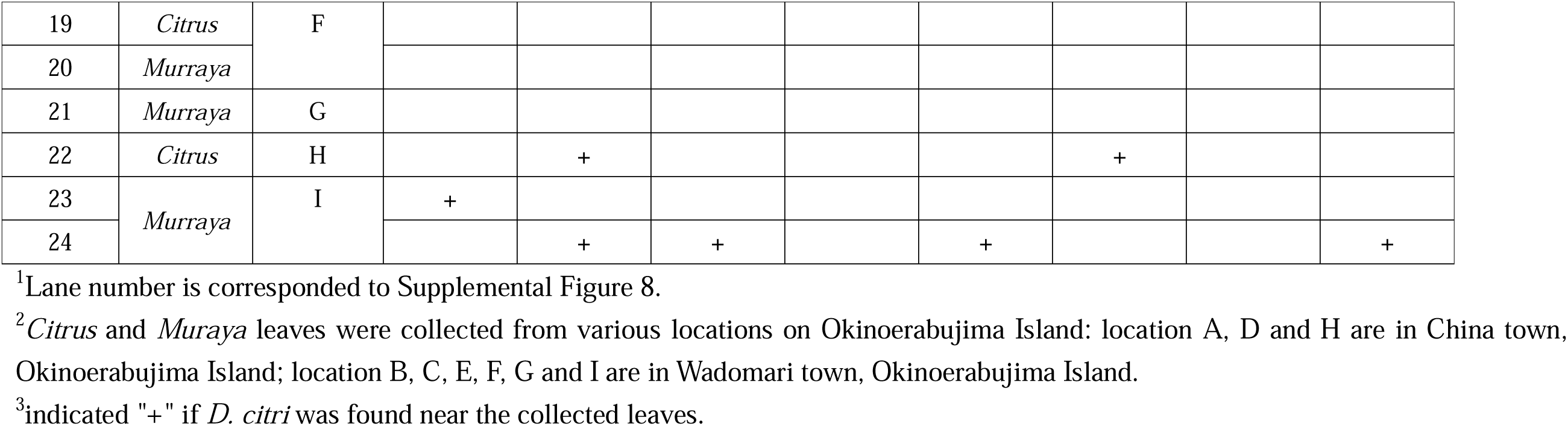
cPCR results using each primer with the DNAs of *Citrus* and *Murraya* leaves in Okinoerabujima Island and presence of *D. citri*.

**Table 4.** qPCR results using each primer with the DNAs of *Citrus* and *Murraya* leaves in Okinoerabujima Island and presence of *D. citri*.

| Samples<sup>1</sup> |  | Dc12S | DcCOI | DcND4 | Wol | wPh | Car | Pro |
| --- | --- | --- | --- | --- | --- | --- | --- | --- |
| *Citrus*<br>location I | 1-1 | –<sup>2</sup> | – | 38. 51 | – | – | – | – |
|  | 1-2 | – | – | – | – | – | – | – |
|  | 1-3 | – | – | – | – | – | – | – |
|  | 1-4 | – | 37. 46 | 36. 91 | – | – | – | – |
|  | 2-1 | – | – | 35. 16 | – | – | – | – |
|  | 2-2 | – | – | – | – | – | – | – |
|  | 2-3 | – | – | – | – | – | – | – |
|  | 2-4 | – | – | – | – | – | – | – |
|  | 3-1 | – | – | – | – | – | – | – |
|  | 3-2 | – | 41. 27 | 34. 80 | – | – | – | – |
|  | 3-3 | – | – | 36. 18 | – | – | – | – |
|  | 4-1 | – | – | – | – | – | – | – |
|  | 4-2 | – | 36. 61 | 36. 23 | – | – | – | – |
|  | 4-3 | – | 42. 02 | – | – | – | – | – |
| *Murraya*<br>location J | 1-1 | – | – | – | – | – | – | – |
|  | 1-2 | – | – | – | – | 35. 73 | – | – |
|  | 1-3 | – | 38. 16 | 35. 53 | – | – | – | – |
|  | 2-1 | – | – | – | – | – | – | – |
|  | 2-2 | – | 39. 53 | – | – | 35. 42 | – | – |
|  | 2-3 | – | 36. 28 | – | – | 35. 37 | – | – |
|  | 2-4 | – | 38. 36 | 36. 91 | – | – | – | – |
|  | 2-5 | – | – | 35. 21 | – | – | – | – |
|  | 2-6 | – | – | – | – | – | – | – |
|  | 3-1 | – | – | 37. 14 | – | – | – | – |
|  | 3-2 | – | – | – | – | – | – | – |
|  | 3-3 | 41. 90 | – | 34. 77 | – | – | – | – |
|  | 4-1 | – | 38. 05 | 37. 26 | – | – | – | – |
|  | 4-2 | – | – | – | – | – | – | – |
|  | 4-3 | – | – | – | – | – | – | – |
|  | 4-4 | – | 36. 89 | – | – | – | – | – |
|  | 4-5 | - | - | - | - | - | - | - |
| *Citrus*<br>location J | 1-1 | - | - | 36.28 | - | - | - | - |
|  | 1-2 | 26.00 | 43.79 | 25.33 | 34.06 | 31.61 | 29.77 | 29.83 |
|  | 3-1 | - | - | - | - | - | - | - |
|  | 3-2 | - | - | 41.64 | - | - | - | - |
| *D. citri* (Positive control) |  | 14.12 | 17.75 | 18.63 | 16.19 | 14.92 | 13.51 | 13.25 |
| water (Negative control) |  | - | - | - | - | - | - | - |

<sup>1</sup> *Citrus* and *Muraya* leaves were collected from Okinoerabujima Island: location I and J are in China town, Okinoerabujima Island.
<sup>2</sup> it was indicated as “-” when the Ct value was not detected.

## Discussion

We succeeded in tracking *D. citri* contact by detecting *D. citri*-derived eDNA from host plants. We believe this is highly innovative as it allows us to monitor the presence of *D. citri* without having to discover the insect itself. Generally, the presence of *D. citri* in the field is confirmed by morphological or DNA-based identification of the collected insect samples (Grootaert et al., 2010; Monzo et al., 2015; Boykin et al., 2012). In the former case, insects were collected in trays by beating host branches or by sweeping the canopy with sweeping nets, or insects were collected with aspirators (e.g., mouth pooter), and then examined morphologically as *D. citri*. Alternatively, yellow sticky traps can be set up in the survey area in advance, and the morphology of the insects caught can be examined (Hall et al., 2010). In the latter case, *D. citri* was identified using PCR or sequencing. PCR primers targeting the mitochondrial COI gene of *D. citri* designed to investigate *D. citri* haplotypes have been used in both PCR and sequencing (Boykin et al., 2012). The advantage of the former is that it is easy to perform; however, its disadvantage is that specialized observation skills are essential. The biggest advantage of the latter is that it has high detection accuracy but the disadvantage is that insects that are likely to be *D. citri* must be screened in advance. Therefore, a method to detect *D. citri* in trap samples using PCR, which combined the advantages of both methods, was developed (Fujiwara et al., 2017). This method is simple and highly accurate because it detects *D. citri* COI gene DNA using bulk samples of insects caught in traps without relying on “artisanal” morphological observations. It is also a very useful method in that it can detect *D. citri* even in samples that have been degraded by UV light or weathering or in samples with only part of the insect body remaining. However, this method requires yellow sticky traps to be set up onsite in advance. Although it can be used to monitor *D. citri* without problems in areas where *D. citri* is constantly occurring and regular surveys are being conducted, it cannot be used immediately in areas where new surveys are required. In particular, in the case of urgent surveys in areas where citrus greening disease has not occurred, setting traps and leaving them may provide time for the disease to spread. In addition, during the initial invasion of *D. citri*, its low population density of *D. citri* makes it difficult to detect by methods such as beating and sweeping. To solve these problems, we devised a method to detect *D. citri* by treating host plants of *D. citri* as traps. Furthermore, we targeted *D. citri*-derived eDNA as a trace of *D. citri*. As targets of *D. citri*-derived eDNA, we attempted to use not only the DNA contained in the mitochondria present in insect cells but also the DNA of symbiotic microorganisms.

It is desirable for detected genes to contain a large number of copies. Therefore, among the DNA derived from insects, the COI gene, which is present in the mitochondrial genome, is often used (Uchida et al., 2020; Yasashimoto et al., 2021; Roger et al., 2021). In addition, genes in the mitochondrial genome, particularly COI genes, are highly conserved and are, therefore, usually used as barcode markers to identify insect species. In this study, we designed primers with high specificity for *D. citri* for the 12S, COI, and ND4 genes, which are conserved in the *D. citri* mitochondrial genome. PCRs using the three primer sets designed based on *D. citri* mitochondrial genes were *D. citri* specific, regardless of whether cPCR or qPCR was used. Particularly, Dc12S and DcCOI effectively distinguished *D. citri* from other Psyllidea and Hemiptera species (Suppl. Table 1; Suppl. Fig 4). However, regarding DNA detection performance, there is a trade-off between the high conservation of target genes in insects and the specificity for detecting only *D. citri*. In fact, by qPCR using DcND4, some Psylloidea were determined to be positive (Suppl. Table 1). Therefore, we designed PCR primers targeting symbiotic microorganisms unique to *D. citri*. Many insects contain bacteria in their bodies (bacteriomes, midguts, salivary glands, etc.) (Brownlie & Johnson, 2009; Ferrari & Vavre, 2011; Nakabachi et al., 2013; Mondal et al., 2023).

Bacterial groups that live symbiotically with insects as hosts are called symbiotic bacteria and are reported to contain host insect-specific bacterial species or form specific bacterial flora. Symbiotic bacteria, such as *Wolbachia, Ca.* Carsonella, and *Ca.* Profftella exist in the bodies of *D. citri* (Saha et al., 2012; Nakabachi et al., 2013; Chu et al., 2016; Hosseinzadeh et al., 2019). Recently, basic and genomic studies on *D. citri*-symbionts interactions have been conducted and remarkable discoveries have been made. However, the development of a detection technology for discovering *D. citri* using the DNA of these symbiotic bacteria has never been envisioned. We read the draft genome of *D. citri* and searched for DNA sequences from symbiotic microorganisms contained therein. We confirmed that all *D. citri* strains used in this study contained the DNA sequences of *Wolbachia*, phage of *Wolbachia*, *Ca.* Carsonella, and *Ca.* Profftella without individual differences. In addition to the three known symbiotic bacteria, we identified *Wolbachia* phage DNA sequences and designed *D. citri* symbiont primers based on these sequences. When we investigated the *D. citri* detection specificity of PCR using these symbiont-specific primers, we found that some insects other than *D. citri* were positive for *Wolbachia* and *Wolbachia* phages but only *D. citri* was positive for both *Ca.* Carsonella and *Ca.* Profftella (Suppl. Table 1; Suppl. Fig 4).

The detection frequency could be increased by preparing multiple *D. citri* detection primers and performing multiple PCRs. We consider it innovative that the test samples were host plants and not *D. citri* itself. Traces of *D. citri* can be detected independently of visual inspection if plant samples contain residual *D. citri* debris (such as parts of the legs or wings) or *D. citri*-derived substances such as eggs, honeydew, and saliva. We confirmed that *D. citri*-derived eDNA can be detected at high frequencies in trace surveys using *Murraya* in experimental greenhouses. Intriguingly, the DNA of *D. citri* symbionts *Ca.* Carsonella and *Ca.* Profftella was detected in the leaves of *D. citri’*s hosts *Murraya* and *Citrus*. This is because, while *Wolbachia* and possibly *Wolbachia* phages are detected in various parts of *D. citri*’s body, *Ca.* Carsonella and *Ca.* Profftella are localized in the bacteriome or female ovaries, making it unlikely that these bacteria were released on the leaves. Bacteriome, where *Ca.* Carsonella and *Ca.* Profftella are primarily localized and specialized organs found in insects of the order Hemiptera that house obligate endosymbionts, where microbial communities are organized to promote insect growth or respond to the environment to improve survival (Nakabachi et al., 2013; Hosseinzadeh et al., 2019). For this reason, even if the remains of the body parts, such as the heads and legs, are on the leaves, it is difficult to imagine that the DNA of *Ca.* Carsonella and *Ca.* Profftella was also detected. In contrast, in the case of eggs, bacteria are supplied by vertical transmission from the female parent; therefore, if eggs (or possibly their remains) are attached to the leaves, DNA will likely be detected. Alternatively, if dead symbionts are released from the body of *D. citri* along with excreta owing to the metabolism of the bacterial flora within the bacteriome, bacterial DNA may be detected on the leaves. In the *D. citri* contact duration tests (Fig. 4; Suppl. Fig. 6), DNA of *Ca.* Carsonella (minimum 10 min) and *Ca.* Profftella (minimum 1 h) was detected even when *D. citri* was brought into contact with the leaves for a short time (no spawning was confirmed), suggesting that DNA traces were left on the leaves due to the excretion of *D. citri*.

If *D. citri*-derived eDNA can only be detected for a short period following *D. citri* contact, the survey period will be limited, making it difficult to use as a method for invasion monitoring and understanding the distribution area of *D. citri*. Therefore, we investigated how long *D. citri*-derived eDNA could be detected in the leaves after contact with *D. citri*. eDNA could be detected even after 180 d, although the detected number of *D. citri*-derived eDNA tended to decrease as the residual period increased (Fig. 5; Suppl. Table 5; Suppl. Table 6). Although this test was carried out in a closed greenhouse without rain or wind, the *D. citri*-derived eDNA remained detectable for up to 180 days (at the end of the test) following contact times of 1 or 7 days. A recent study found that although the residual amount of *D. citri* DNA on sticky traps in practical fields in Florida varied depending on the season and the amount of amplification varied depending on the type of target gene in PCR, it was still detected even after four weeks (Bloch et al., 2023). Thus, *D. citri*-derived eDNA may remain undegraded on leaves for longer than expected.

Therefore, we tested whether our newly developed method for detecting traces of *D. citri* using eDNA could be used in practical *D. citri* habitats. On Okinoerabujima Island, in the southwest archipelago of Japan, Murraya and Citrus trees grow indigenous to the mountains or are planted in home gardens. *Diaphorina citri* inhabits and citrus greening disease occurs here; therefore, ongoing control efforts are being carried out, such as monitoring this disease and cutting down diseased trees as early as possible to prevent the spread of the disease via *D. citri* (Iwanami, 2022). Vector control is also progressing, and the *D. citri* population is decreasing. Detecting *D. citri* in these areas is difficult using conventional methods owing to seasonal restrictions as well as various constraints such as personnel, labor hours, and expenses. Therefore, when we investigated whether our new method of monitoring *D. citri* habitats by detecting eDNA could be used in a practical survey, we detected *D. citri*-derived eDNA in many leaves on site (Table 3; Table 4). Although it cannot be determined whether these are true traces of *D. citri*, using primer sets that can distinguish *D. citri* from the closely related insects *Psylloidae* and *Hempitera*, and using primer sets that detect *Ca.* Carsonella and *Ca.* Profftella are *D. citri*-specific symbiotic bacteria, and positive PCR results were obtained from the DNA of leaves at the site (Suppl. Table 1; Suppl. Fig. 4). Therefore, it can be concluded that these compounds were derived from *D. citri*. In fact, *D. citri* individuals were sometimes found near the leaves where eDNA was detected. Therefore, this method may be useful in areas where *D. citri* has not occurred or where its population is extremely small, when it is desirable to quickly find traces of *D. citri*. Traces of *D. citri* DNA remaining on leaves cannot always be detected by PCR using each primer set because they are completely or partially degraded by various factors in the field. However, when a low frequency or a few (sometimes single) PCR results are positive, *D. citri* may be found by surveying trees and surrounding host plants. Furthermore, it would also be possible to simultaneously investigate whether host plants are affected by citrus greening. During the field applicability test of this method, highly accurate PCR for detecting citrus greening disease bacteria (Fujikawa & Iwanami, 2012; Fujikawa et al., 2013) was performed on the sample DNA but no positive results were obtained (data not shown). To implement this method for *D. citri* on-site monitoring, evaluation of various abilities, such as efficiency, sensitivity, accuracy, and frequency, using field host plants and technical improvements are essential. In the future, we believe that our method of detecting *D. citri*-derived eDNA in leaves to find traces of *D. citri* will enable rapid monitoring of *D. citri*, a vector of citrus greening disease. This will prevent the invasion and spread of citrus greening diseases and contribute to the stabilization of citrus production worldwide.

## Author Contributions

TF, KF, and HI conceived and designed the study. TF, KF, HI, HH, and SH performed the experiments. TF contributed to the data analysis. TF, KF, and HI contributed to the manuscript writing.

## Acknowledgments

We thank the members of the Institute for Plant Protection, NARO, for helpful discussions. We would like to thank Editage (www.editage.jp) for the English language editing.

