## Supplementary Text 1 for "Investigation of host plant contact with *Diaphorina citri* (Hemipreta: Psyllidae) by detecting *D. citri*-derived environmental DNA": 11.Suppl. text 1 contig seq-ed1.docx

**Supplemental Text 1**

#1 Contig of Wolbachia spp.

Contig_136

2,269 bp

Predicted genes: Rpn family recombination-promoting nuclease/putative transposase

>Contig_136_Wolbachia_pipentis

TTTGGATGCCTTCTTGTCTACCTTCTTGTCTACCTTCTTGTCTATCTTCTTGTTTAGTCATCATCGAGTTTTTGAGCAAGGATAGCCTTCTCTTTCTGCACACTCAATATTCTTTCCTCATATGCGATCAAATCTTTTTCATTCCAGCGAAACCTGTCTAATTCATCATATGCTAGCTTTATTATCGGAGCTTCTGCTGCTATCTTTTTCAGATCTTCTTCAGTTGTATCTTCTGCGTATTTAAAAAAGAAACACCATCTCTCTGTAGTATTCTCTAACTGCTCTACTCTATTTTTTGTAAATTTAGGCAACTCAATAAAGACAAATTGTAGATCTTTCAAGTAATGCCCATTGGTTTTGATATCGCGTATATTATGAGTAGAAATATAGTCAACTTCTTCAGGAAGAAGGTTACAATTGGAAATAGCAATAAAGAAGACTTTCTGAAGATCAATGTAATTACCAGATTTATCTAATTGTCTTGAGTAAGCCTTAGCAACATAAAGTTGAGCACGTTTTTCAAAGCCTTTATCACGAGCAAGCTGCATTTCCGCAATATACCTATTTCCGAGAGAATCCTTACAGAGGACATCAACAATGCTTTGTTTATCAGAAGCAATCTCAGGATTCATAATGGTGCTGAGAAACTCAACATCCTGAATAGCATTAACTCCAGTAAAGCCTAAGATATCATTCAAAAAATGGATAAGGATATTCTTGTTTTTTTCAGTACCGAAGATTTTCTTAAATGTTAAATCTAATTTTGGATCGAGAAACTTCGAAAGAGCCATGAGAAAGTAAGATAAAAAGCATTAATAATTATACACAATTGTGAAGAAATATTCAACTAAAATTCAAACATAGAAAGAAGCATCCCTTTTGTATATCTTTATTTGGTAGTATTAACCTGTATAGTATTTAAGAACATTAGCCCTGCAAGATACAAAATAGCACAAAAAATAAGAAGCTGGAGGCTAAAACAAAAGTATACCCTAAAAGATCTAGTAGAGAAAACAGGTATAAAGTATCATATGTTGCTAAGATATGAGAAAGGAACATGTGGCATTCCAATTGAGAAGTTAAGAATATTAGCAGATGCATTATCAATCCTATTAGAGATCTCTTTCCAAGACGAAAAGTACTAAAAGAAAGCAGTTGTTTTGACAAAGCCAAGACCCAGGAAATGTATAATTCATAGAAAAAAACAGGAGGACACAAAGCAATTTATATGTTAACCAAATCTGTCCGAGCTGAGGAAGAAAGTAATATAAAAGCAGCAAGAATAAGAATTGCAAGGAATCTAGTTAAGGCAGAGTTTGATACTGACATTATCTATCGAACAACAGGCTTATCAACTGAAGAATATGCTGATAAAGAGAGAGAGCATTCAAACGAAGGACAAGAGATAAAAAAGTGGAGGATAATAAGAGGGTATACTCAAGAGGATTTAGCAAGAGAGCTAGACATAGGACCTTCACAGATACATCAACTATGAACAAGGTAGTGTTACTATTTTAAGTGAAAGGTTATGGGAAATAGCAAAAGAATTATCAGTGAATGCTGAAGATCTGATAAAGGAGTACAAAGAAAATGATTGCGAAGGAGAAAGTGAATTATTAAGTTTGGCAAGAGAATATAGAAAAATTGACAATCAAGAATCACGAGATGAGCTAAATATATGGGTAGAATTGTTATTGCAAAGAAAGCAAATTTACAAAGAAAAGATTTACAAAACAGAGGAAATGAAAGTTGCAAATAATTTACTTCAGTTAGGTTTTTCTACTGATGTTATTTCTAAAATAGTAACAACTTCTTTTACATAAAGAATTAATTTTCAATTAGATAAAATAAGCATACGTTAATATCTTAAATTTTTGATAAAAAAATAAGTAAAAAAGATTGAAAAATTGATTTGACTTCACTCGGAAGCTATGGCTCTATGCCAATGCTGATAAAAAACAGCAGTAAATAATTAAAAACAATTTGTTATAAGAAAAAAAAAAAATGTAATAAATGGAAAAAAAAAAAAGAAAAAAAGAGGGGATAGGGGGTATGTATATCTGTAACAACACAGGGCTTTTTATCCCGAAATCGAAGAGAAAATCAGGACAGGAAGGCATTACTTACCTAAGCTTTCTATTAAGGGGCAGTGGCACCATTTTCCTTGAAGGAGCTCATCCTTCAATGAGTAGCTTAAACAATCAGATCCCCCATTCTCTTTCTAGTCGAATGCTTAATCCAAATGGATTAAGAATGCTTAATCCAAATG

#2 Contig of Wolbachia phage

Contig_148

1,107 bp

Predicted genes: DUF106 domain-containing protein

>Contig_148_Wolbachia_phage

TGTTGTGCGGTACGGGGAAGTTCAAATAAAGGATTTATTGGAGCTCAGGCAACGAATAAAAGCGGGGTTAAAAGTTGCAGGCATGAGGCCAAAGAGGAAAATTGTTTTTTCAACGAGTAAGGGGATCATATAATGATAAAAATCGTTAAAAGGAACCGTATGACAAAAATTATAGCAAAAGAATATACAGAATTTTTAGAGCAGCTCAAAGAACAGATTGCTACTAGTCGTTATAAAGCAGCACTAGCAGTAAATAGCAAGCTTATTGTGCTTTACCACCATATTGGTACAGAGATCTTAAAACGCCAAAAAGAGCATGGCTGGGGTGCTAAAATCATAGATCAGCTAAGTAAGGATTTGAGAAGCGCATTTCCAGAAATGAAGGGATTTAGTGCTAGGAATCTTACTTATATGCGTCAATTTGCTGCAGAATATCAAGATACTGAATTTACGCAGCAGGTTGCTGCACAATTGCCTTGGTTTCATATTGTAGTTATTTTAGATAAAGTAAAAGATCCAAAAGAAAGACTATTTTATTTTCAAAAAGATATAGAGCACGGCTGGTCACGTAGTATAATGGTGATGCAAATCGAACGTGAATTGCACAAACGTCAAGGAAAAGCTGTTACTAACTTCAAAGATAAGTTACCTTCTCCCCAGTCAGATTTAGCACATTATACATTTAAAAGATCCATATATTTTGATTTTTTGAGTATAGGAGATGACGCTCATGAAAGAGAAGTAGAAAAAGGTCTTGTAGGCCATGTGGAAAAATTTCTGCTTGAGCTAGGAGAAGGATTTGCATTTGTAGGTAGGCAATTTCACTTAGATGTTGGAAATAAAGACTTTTATATTGATTTATTGTTTTACCATTTAAAGTTACGTTGCTTTGTAGTAATTGAGCTAAAAGACAAAGATTTCAAGCCAGAATATGCAGGTAAAATGAATTTTTATCTTTCAGCCGTTGATGATTTACTAAAGCACAAGACAGATCAACCATCTATAGGATTAATTTTATGCAAGTCAAAGGATAACGTGCTAGCTAAGTATACACTGAAAGACATAAATAAACCGATAGGGTTGGCAGAATATAAAATTACTGAAAAT

#3 Contig of *Candidatus* Carsonella spp.

Contig_207

1,217 bp

Predicted genes: Carsonella 23 S rDNA

>Contig_207_Carsonella

CACAAAATAATGTTCATAATTTCGGTACTTATTTTAGCCCCGTTAAATTTTTCGAATTAATCATCTATATCAATGAGCTATTACGCTTTCTTTAAAGGATGGCTGCTTCTAAGCCCACCTTTTGATTGTCAAAGATTATTAAAATTCATTTTCCACTTAAATAAGATTTAGGGACCTTAATTTATGATCTGGGTTGTTTCCCTTTTCACAATGGATGTTAGCACCCATTGTGTGTTTCTTATAATAAAAAAACAAATATTCATAGTTTGTTATGATTCAGATAAACTTAATCAAATACAGTGCTTTACCATTAGTTTTCACTTATAAGACGCTACCTAAATAGCTTTCGAGGAGAACCAGCTATCTCCGAGCTTGATTAGCCTTTCACCCCTATCCACAAATCATCTGAATCTTTTGCAACAGATACCAGTTCGGTCCTCCAGTAATGATTATATTACCTTCAACCTGTTCATGGATAGATCGCTCGGTTTCGGGTCTATTATTTTTAACTATCGCTCTTTCAAACTTGATTTCTCTACGCCTACTAATAATTAAGCTTGCTAAAAATAATAAGTCGCTGACCCATTATACAAAAGGTATATAGTTGCTTTTCAGCTTCTATTGCTTTTACGTATATAATTTTAGGTTCTATTTCACTCCTATAAAAAGGTTCTATTTCATCTTTCCCTCACGGTACTAGTTCACTATCGGATAACTATTAGTATTTAGCCTTAGAGGATGGTCCCCCTATATTCTGTTAAGATACTACGTGTCTAAACATACTCATAATAAAAATATAAAACAAAAAATAAAAAAAGACTATTACTTTTTTATGTTAGTTATTCAAAACTATATTATTTTCGTTTAAATATATATTTTAAGCTTCTCCCATTTCGCTCGCCACTACTGTGGGAATCTCATTTGATTTCTTTTCCTTGGATTACTTAGATGTTTCAGTTCATCCAGTTTGCGTTTTACACTAATTAATATTAGTAGGTTACCCCATTAAGATACCTTTTACAAAGATTATCGCATTTTAGCGTCTGTCATCGCCTATAGTTACCAAGACATCCTTTATATACGTTATTTTTTTTAATTTATAAATTAATTTTTATATCAAAATCAAAAATAAGAAATATATTATTATATCCAGCCACAGGTTCCCCTACAGCCTACTTGTTACGACTTCACCCCAGTTATAAATCAATAC CGTCGCA

#4 Contig of *Candidatus* Profftella spp.

Contig_27

818 bp

Predicted genes: Ribonuclease

>Contig_27_Profftella

AAAATAAAATTAAAAAATTATGACCGTAAATGTACTCGCTAAAAAGCGCACCTACCTATCCTACCAATAACTTTAAATACTTTAAAATTTTAAAAAAATATATATTTTATCAAAAATTGTTGATTTAATTGAACTAATTATTTTATCGATTTTACAATTATTTGAATTATAAATTGATTATAAATTTATATTCoTATTTAATAATAACGGAAATAGATTATGAATTTACTTTTGAAAAATATGGTACCTTTAAAGTTGGTACAATTTTTTCTCAGAATGGTAATTTTTATCAAGTTAAAATAATTTATAATTAAATTAAATAATGAGAGCAATATGATATTTTGTTACGATTTTCCACCCCAAATCCAGATCAATTAATTAAAAAAGCAAAACATTATTAGCTAATGAAATTGATCTTAATTTTTTATGGGAAATTGTTGGAACAAAGAAATTTTACTTTTTTGATCTAGGTATAGAATATTTCGGGCATACTCCATTATCATATGAGGCAGCTAGTTTATTATTTTGTTTACATAATTCTCCTATTTTATTTTTATAAAAAAAGGGAAAAGGATATTTATAAAGCGGTACCTAGAGAATTTATTAAAAGACAACTAAAAATTCATTTAGAAAAAAAAAAAATTCAAAAAATTGAAATAATTATGATTAAATTAGTAAATTAAAAAAATAATAATTAAGAATTCGGCAAATTTAATTTTCGACTGTTTATCAAAAACATTTCTGGAGAAAAAATATTTTTGGTATGTCCTGCTCAATGCTGATGTAAATAGCTGCAGTATACTGACTGTACAAAGGTAGC
