## Supplementary Table 3 for "Investigation of host plant contact with *Diaphorina citri* (Hemipreta: Psyllidae) by detecting *D. citri*-derived environmental DNA": 14. Suppl. Table 3.docx

**Supplemental Table 3. Short sequence reads obtained from ten individual *D. citri*.**

| **Individual No.** | **Number of reads** | **Average length (bp)** | **Accession No.** |
| --- | --- | --- | --- |
| 1 | 59,973 | 285.5 | [SRR28552066](https://www.ncbi.nlm.nih.gov/sra/?term=SRR28552066) |
| 2 | 510,686 | 287.4 | [SRR28552065](https://www.ncbi.nlm.nih.gov/sra/?term=SRR28552065) |
| 3 | 593,819 | 276.9 | [SRR28552064](https://www.ncbi.nlm.nih.gov/sra/?term=SRR28552064) |
| 4 | 420,869 | 282.8 | [SRR28552063](https://www.ncbi.nlm.nih.gov/sra/?term=SRR28552063) |
| 5 | 420,869 | 282.8 | [SRR28552062](https://www.ncbi.nlm.nih.gov/sra/?term=SRR28552062) |
| 6 | 1,032,434 | 281.7 | [SRR28552061](https://www.ncbi.nlm.nih.gov/sra/?term=SRR28552061) |
| 7 | 1,290,665 | 265.9 | [SRR28552060](https://www.ncbi.nlm.nih.gov/sra/?term=SRR28552060) |
| 8 | 1,290,665 | 265.9 | [SRR28552059](https://www.ncbi.nlm.nih.gov/sra/?term=SRR28552059) |
| 9 | 683,737 | 278.3 | [SRR28552058](https://www.ncbi.nlm.nih.gov/sra/?term=SRR28552058) |
| 10 | 769,064 | 285.8 | [SRR28552057](https://www.ncbi.nlm.nih.gov/sra/?term=SRR28552057) |
