## Supplementary Figure 1 for "Investigation of host plant contact with *Diaphorina citri* (Hemipreta: Psyllidae) by detecting *D. citri*-derived environmental DNA": 18. Suppl. Figure 1.pptx

### Slide 1
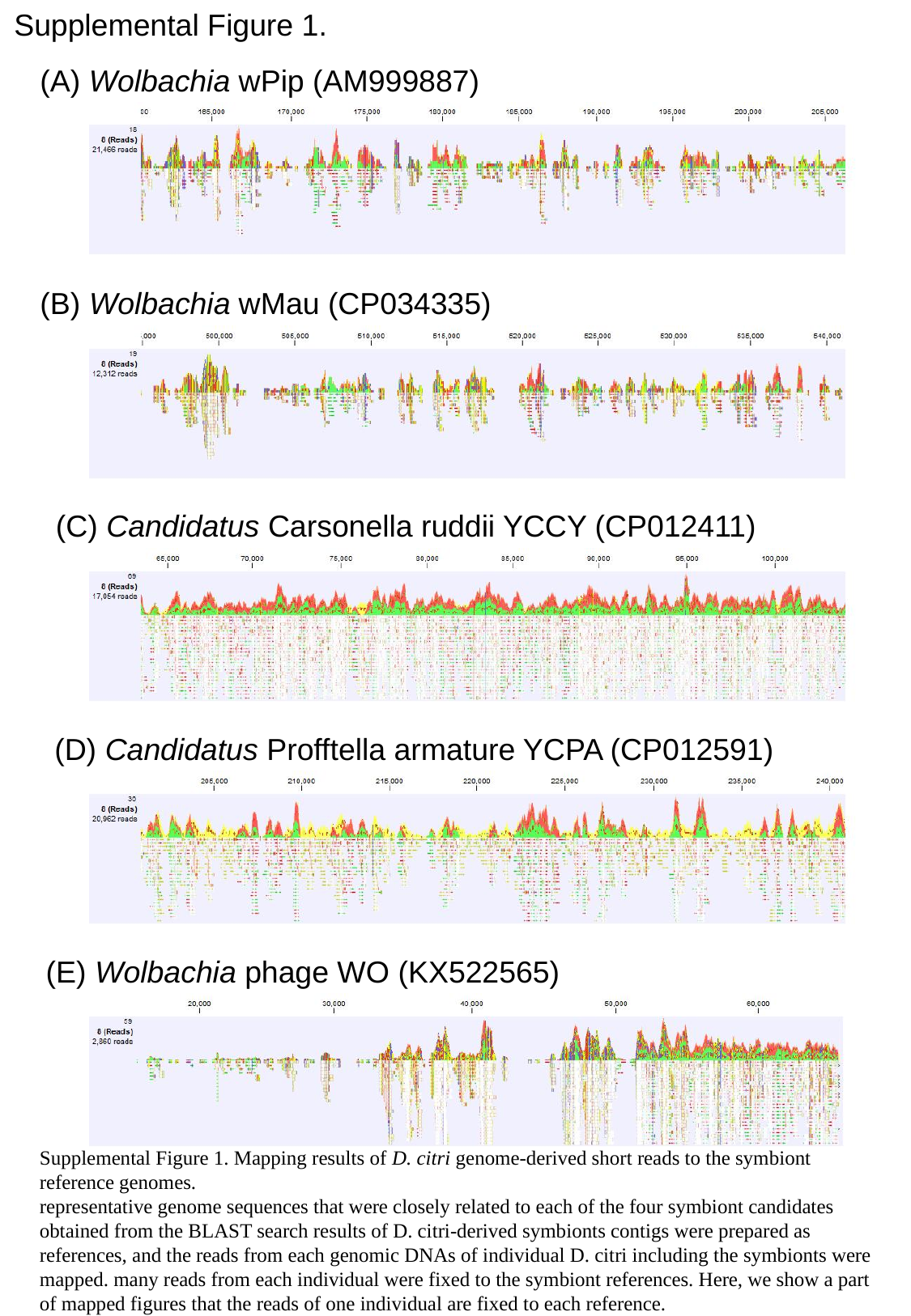

Supplemental Figure 1.
(A) Wolbachia wPip (AM999887)
(B) Wolbachia wMau (CP034335)
(C) Candidatus Carsonella ruddii YCCY (CP012411)
(D) Candidatus Profftella armature YCPA (CP012591)
(E) Wolbachia phage WO (KX522565)
Supplemental Figure 1. Mapping results of D. citri genome-derived short reads to the symbiont reference genomes.
representative genome sequences that were closely related to each of the four symbiont candidates obtained from the BLAST search results of D. citri-derived symbionts contigs were prepared as references, and the reads from each genomic DNAs of individual D. citri including the symbionts were mapped. many reads from each individual were fixed to the symbiont references. Here, we show a part of mapped figures that the reads of one individual are fixed to each reference.
