## Supplementary Figure 2 for "Investigation of host plant contact with *Diaphorina citri* (Hemipreta: Psyllidae) by detecting *D. citri*-derived environmental DNA": 19. Suppl. Figure 2 adult nymph plant.pptx

### Slide 1
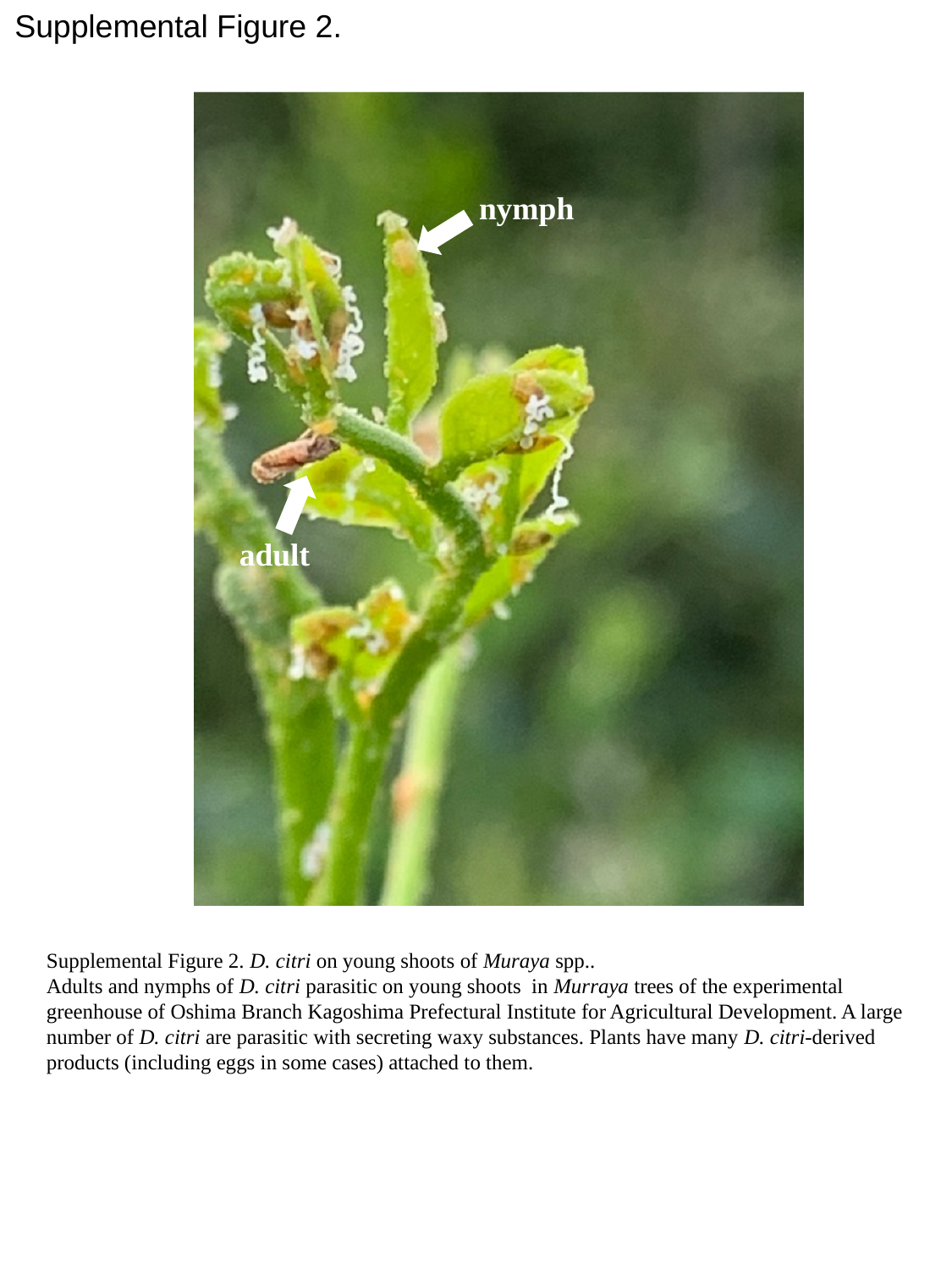

Supplemental Figure 2.
nymph
adult
Supplemental Figure 2. D. citri on young shoots of Muraya spp..
Adults and nymphs of D. citri parasitic on young shoots in Murraya trees of the experimental greenhouse of Oshima Branch Kagoshima Prefectural Institute for Agricultural Development. A large number of D. citri are parasitic with secreting waxy substances. Plants have many D. citri-derived products (including eggs in some cases) attached to them.
