## Supplementary Figure 3 for "Investigation of host plant contact with *Diaphorina citri* (Hemipreta: Psyllidae) by detecting *D. citri*-derived environmental DNA": 20. Suppl. Figure 3 Murraya.pptx

### Slide 1
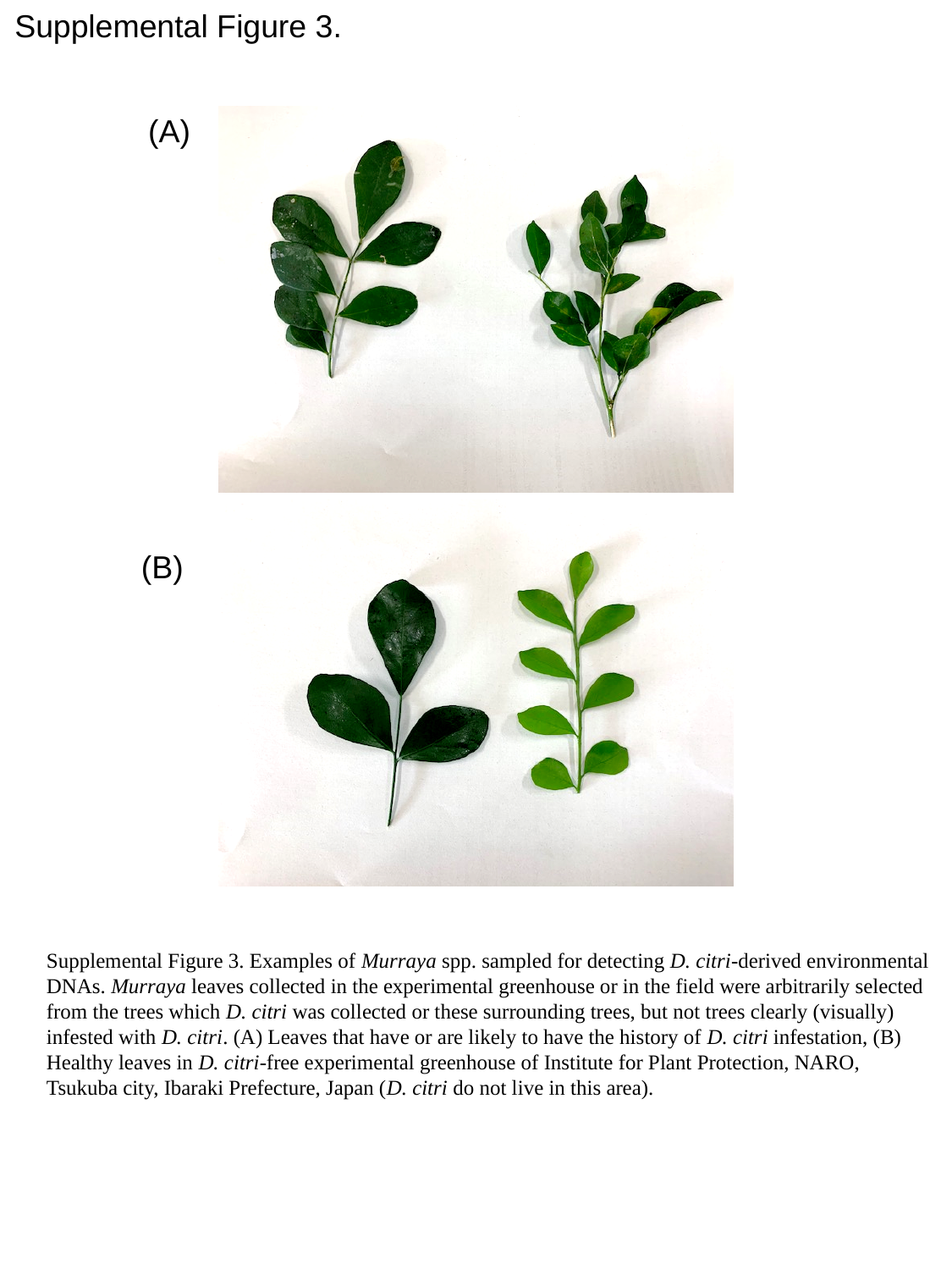

Supplemental Figure 3.
(A)
(B)
Supplemental Figure 3. Examples of Murraya spp. sampled for detecting D. citri-derived environmental DNAs. Murraya leaves collected in the experimental greenhouse or in the field were arbitrarily selected from the trees which D. citri was collected or these surrounding trees, but not trees clearly (visually) infested with D. citri. (A) Leaves that have or are likely to have the history of D. citri infestation, (B) Healthy leaves in D. citri-free experimental greenhouse of Institute for Plant Protection, NARO, Tsukuba city, Ibaraki Prefecture, Japan (D. citri do not live in this area).
