## Supplementary Figure 4 for "Investigation of host plant contact with *Diaphorina citri* (Hemipreta: Psyllidae) by detecting *D. citri*-derived environmental DNA": 21. Suppl. Figure 4 Hemptera.pptx

### Slide 1
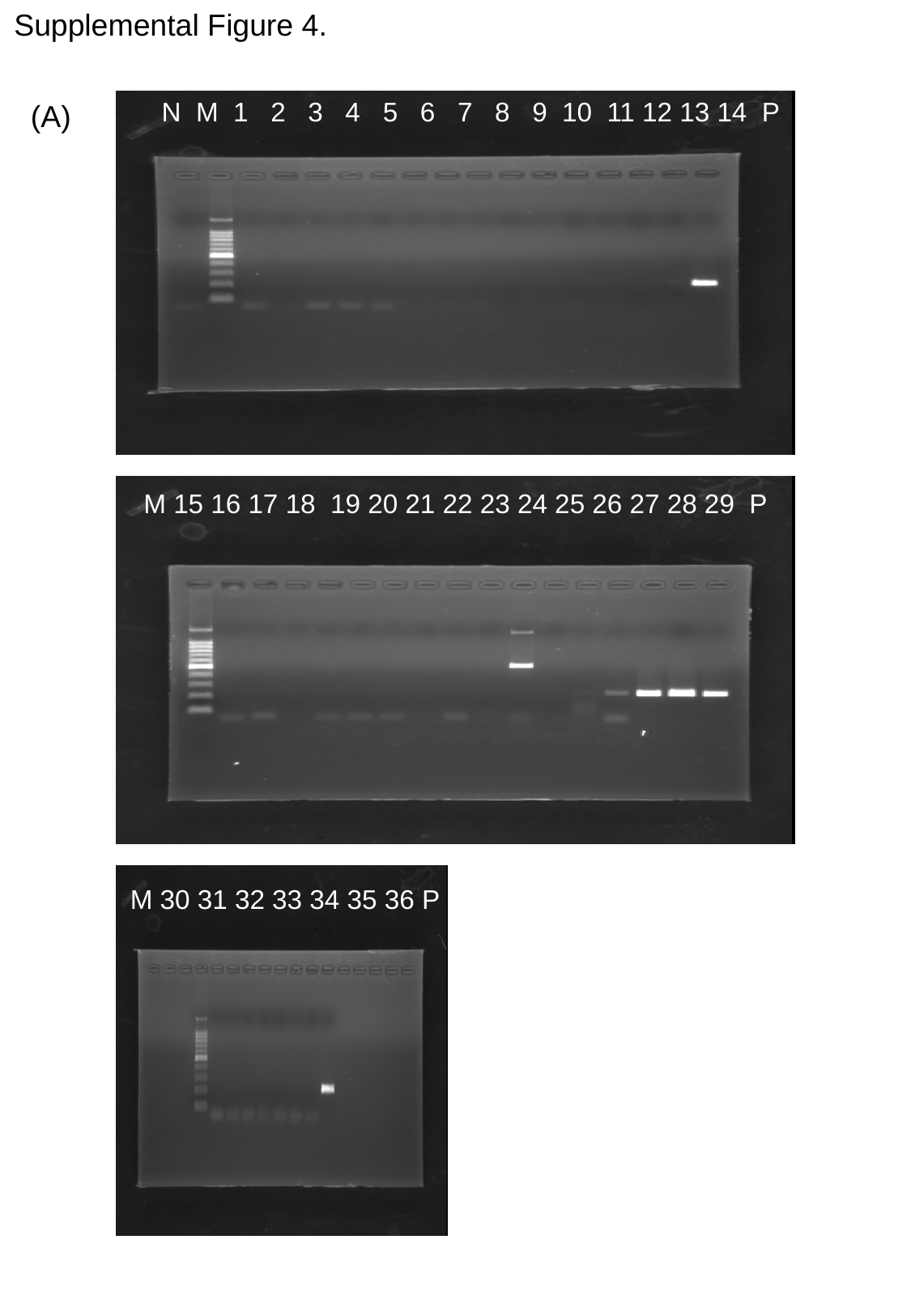

Supplemental Figure 4.
N M 1 2 3 4 5 6 7 8 9 10 11 12 13 14 P
(A)
M 15 16 17 18 19 20 21 22 23 24 25 26 27 28 29 P
M 30 31 32 33 34 35 36 P

### Slide 2
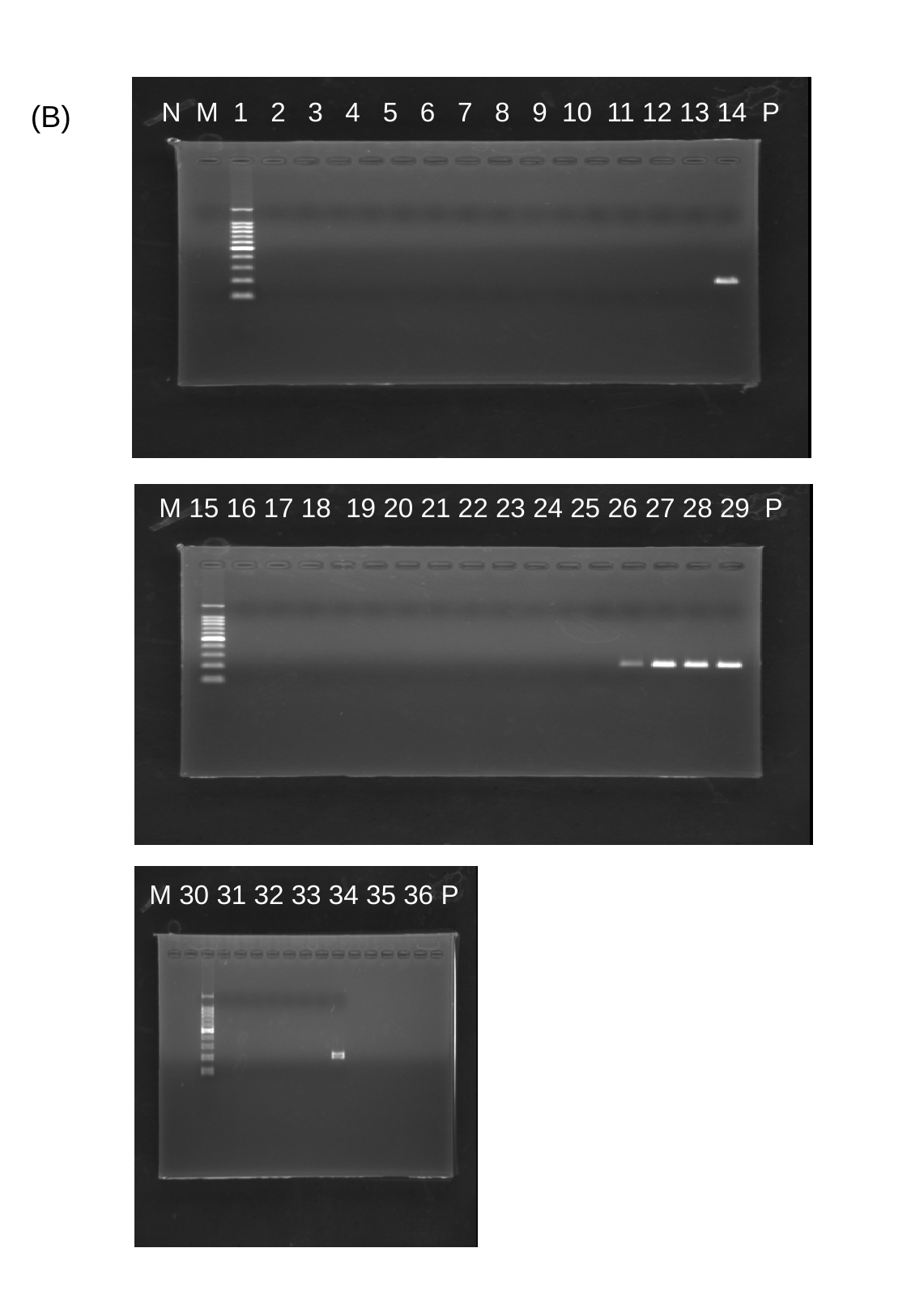

N M 1 2 3 4 5 6 7 8 9 10 11 12 13 14 P
(B)
M 15 16 17 18 19 20 21 22 23 24 25 26 27 28 29 P
M 30 31 32 33 34 35 36 P

### Slide 3
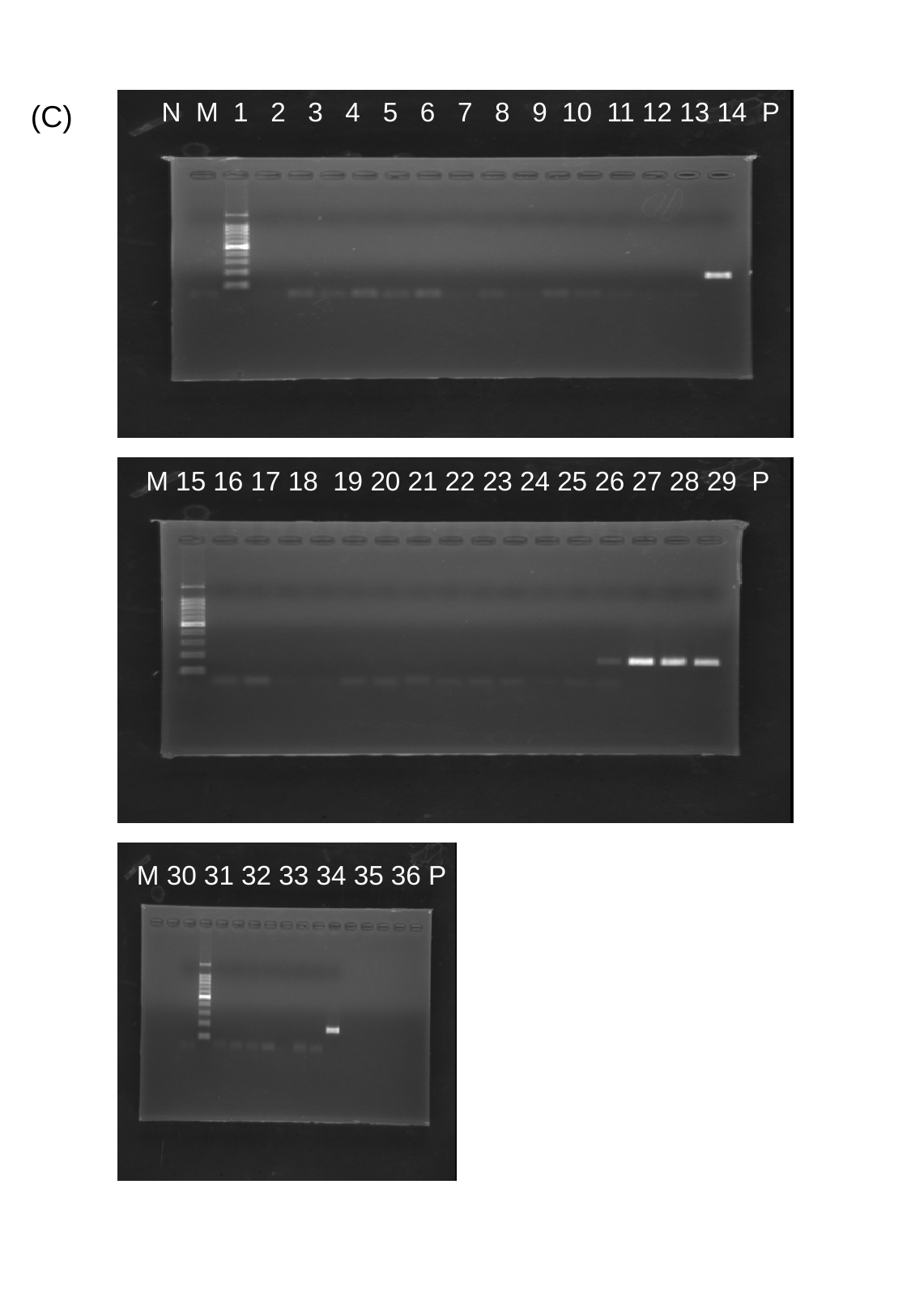

N M 1 2 3 4 5 6 7 8 9 10 11 12 13 14 P
(C)
M 15 16 17 18 19 20 21 22 23 24 25 26 27 28 29 P
M 30 31 32 33 34 35 36 P

### Slide 4
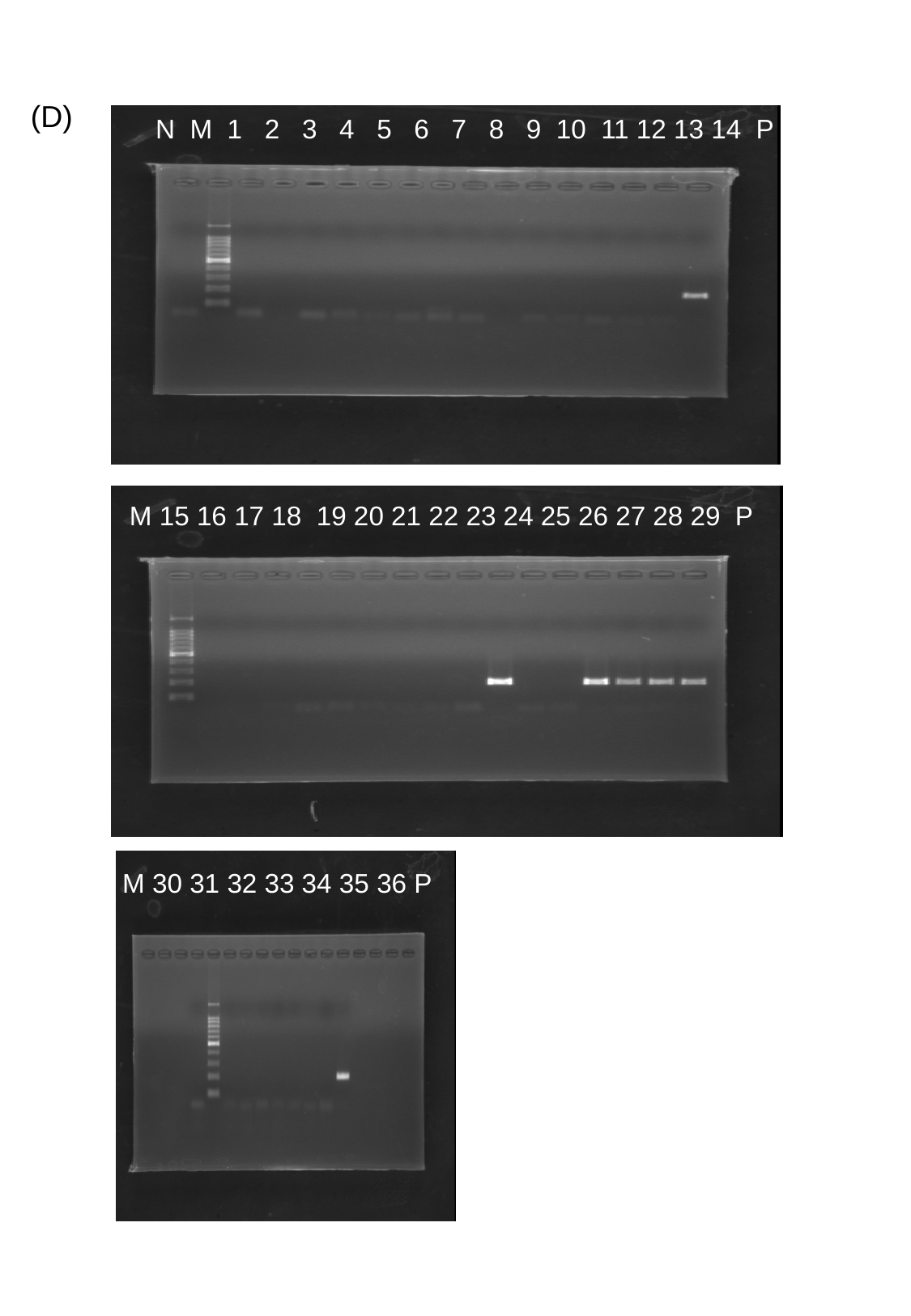

(D)
N M 1 2 3 4 5 6 7 8 9 10 11 12 13 14 P
M 15 16 17 18 19 20 21 22 23 24 25 26 27 28 29 P
M 30 31 32 33 34 35 36 P

### Slide 5
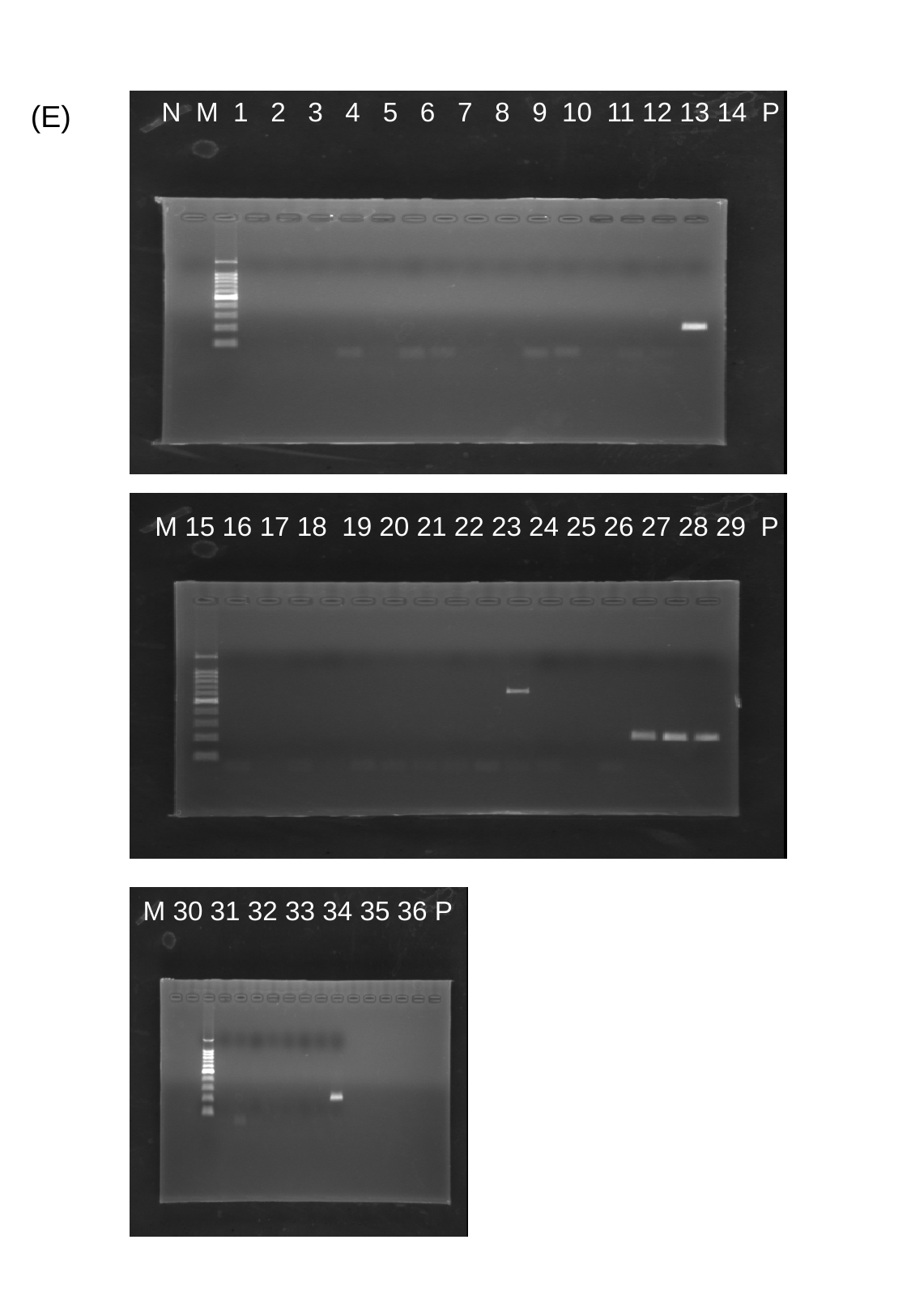

N M 1 2 3 4 5 6 7 8 9 10 11 12 13 14 P
(E)
M 15 16 17 18 19 20 21 22 23 24 25 26 27 28 29 P
M 30 31 32 33 34 35 36 P

### Slide 6
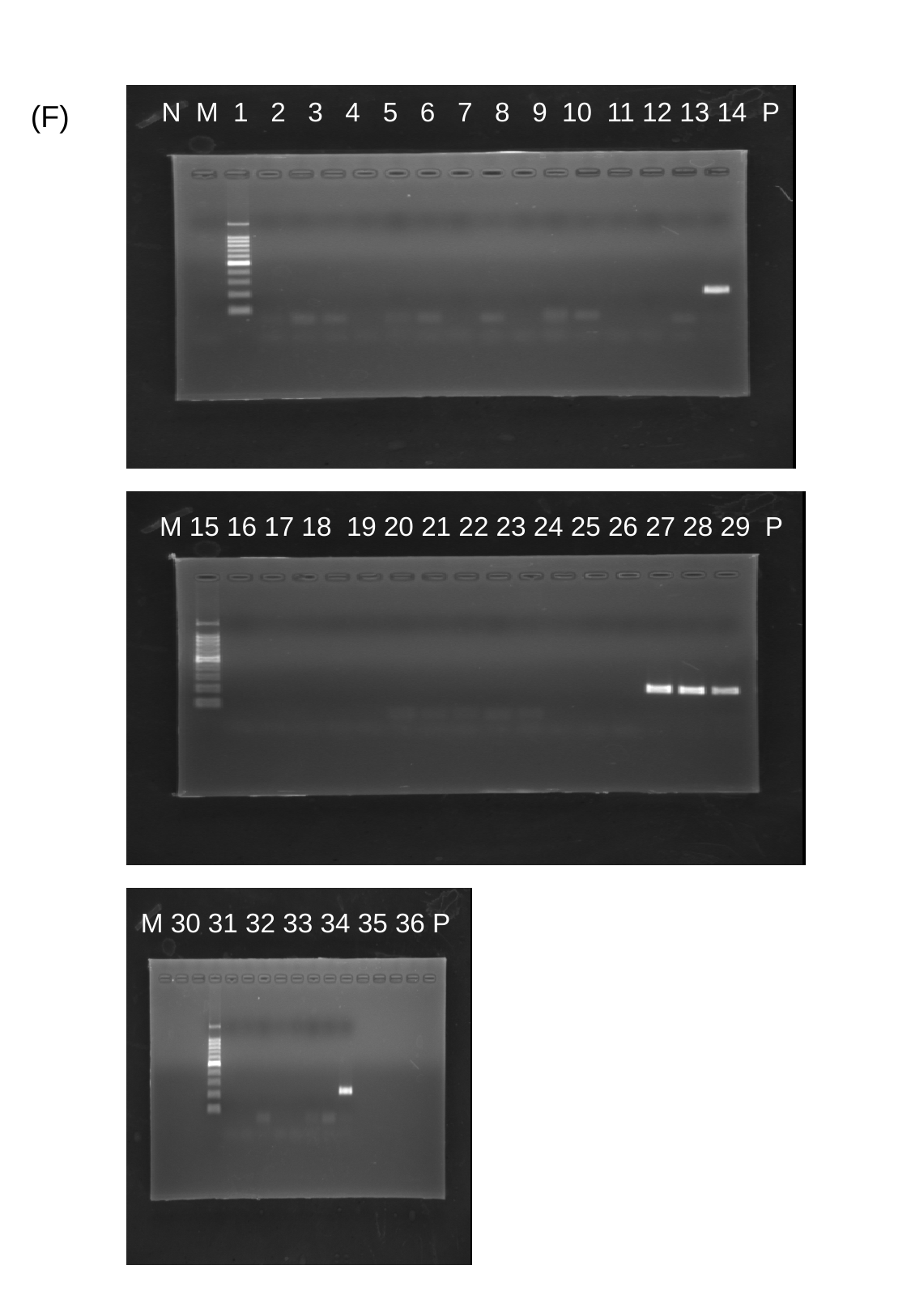

N M 1 2 3 4 5 6 7 8 9 10 11 12 13 14 P
(F)
M 15 16 17 18 19 20 21 22 23 24 25 26 27 28 29 P
M 30 31 32 33 34 35 36 P

### Slide 7
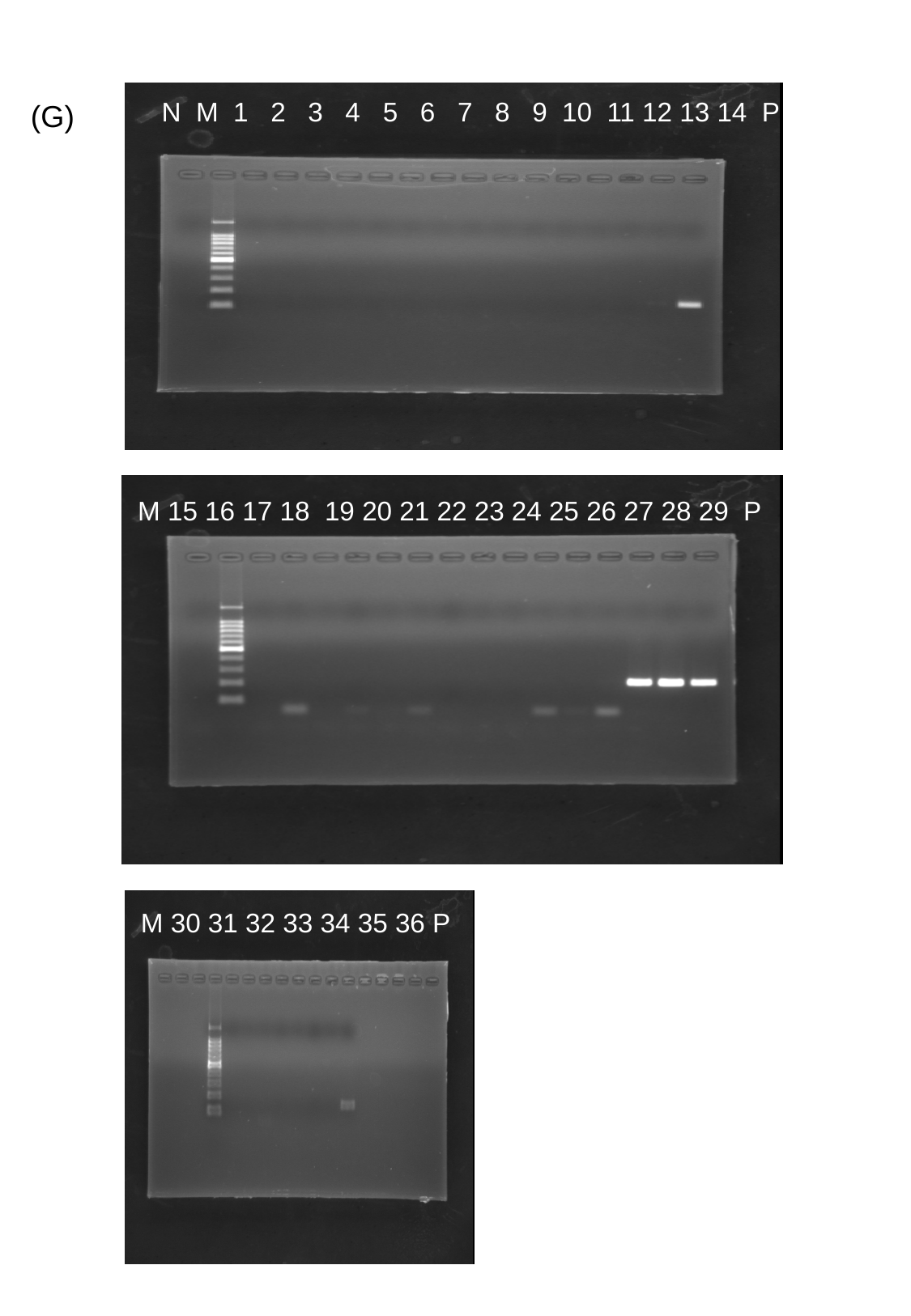

N M 1 2 3 4 5 6 7 8 9 10 11 12 13 14 P
(G)
M 15 16 17 18 19 20 21 22 23 24 25 26 27 28 29 P
M 30 31 32 33 34 35 36 P

### Slide 8
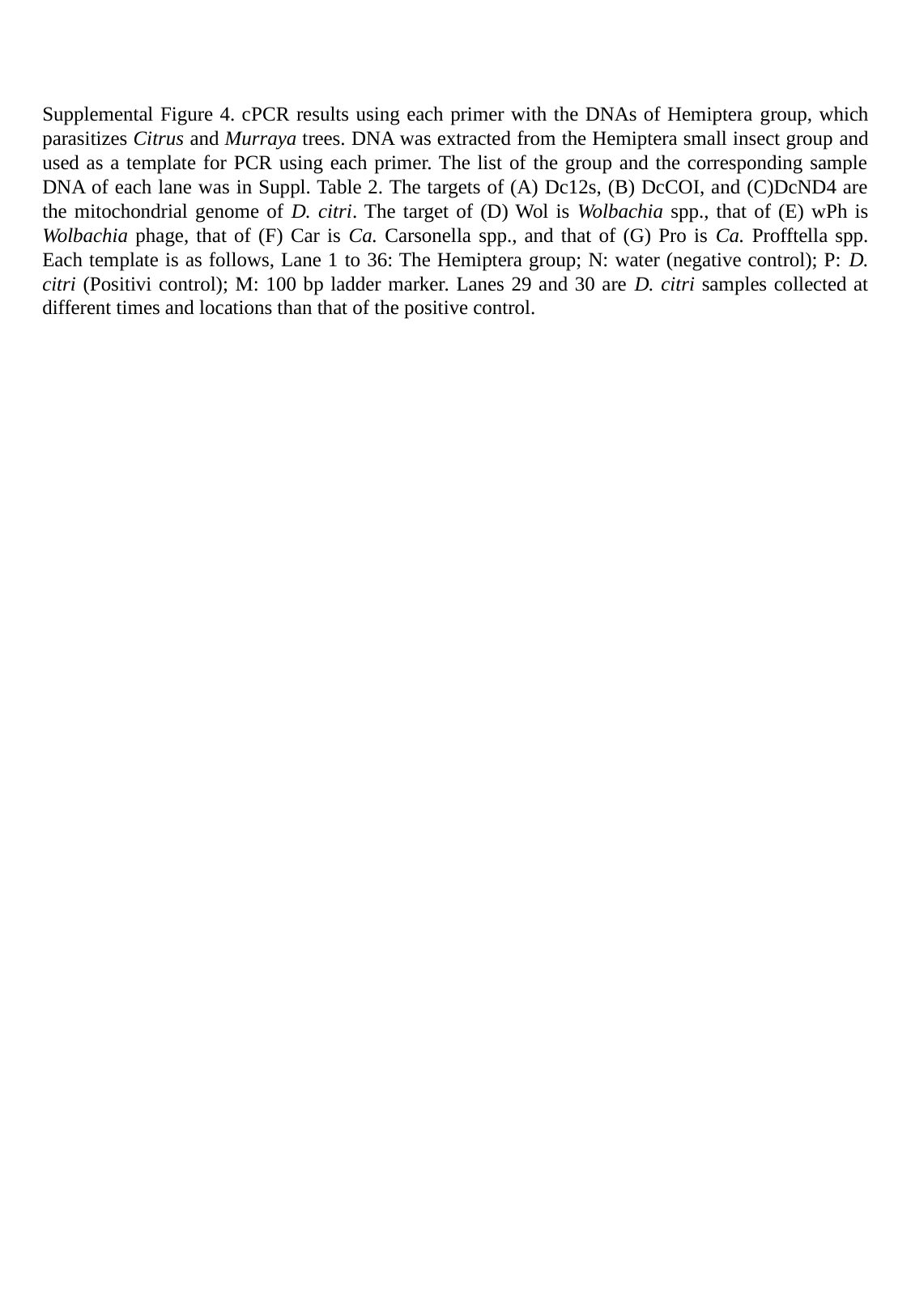

Supplemental Figure 4. cPCR results using each primer with the DNAs of Hemiptera group, which parasitizes Citrus and Murraya trees. DNA was extracted from the Hemiptera small insect group and used as a template for PCR using each primer. The list of the group and the corresponding sample DNA of each lane was in Suppl. Table 2. The targets of (A) Dc12s, (B) DcCOI, and (C)DcND4 are the mitochondrial genome of D. citri. The target of (D) Wol is Wolbachia spp., that of (E) wPh is Wolbachia phage, that of (F) Car is Ca. Carsonella spp., and that of (G) Pro is Ca. Profftella spp. Each template is as follows, Lane 1 to 36: The Hemiptera group; N: water (negative control); P: D. citri (Positivi control); M: 100 bp ladder marker. Lanes 29 and 30 are D. citri samples collected at different times and locations than that of the positive control.
