## Supplementary Figure 5 for "Investigation of host plant contact with *Diaphorina citri* (Hemipreta: Psyllidae) by detecting *D. citri*-derived environmental DNA": 22. Suppl. Figure 5 duration scheme.pptx

### Slide 1
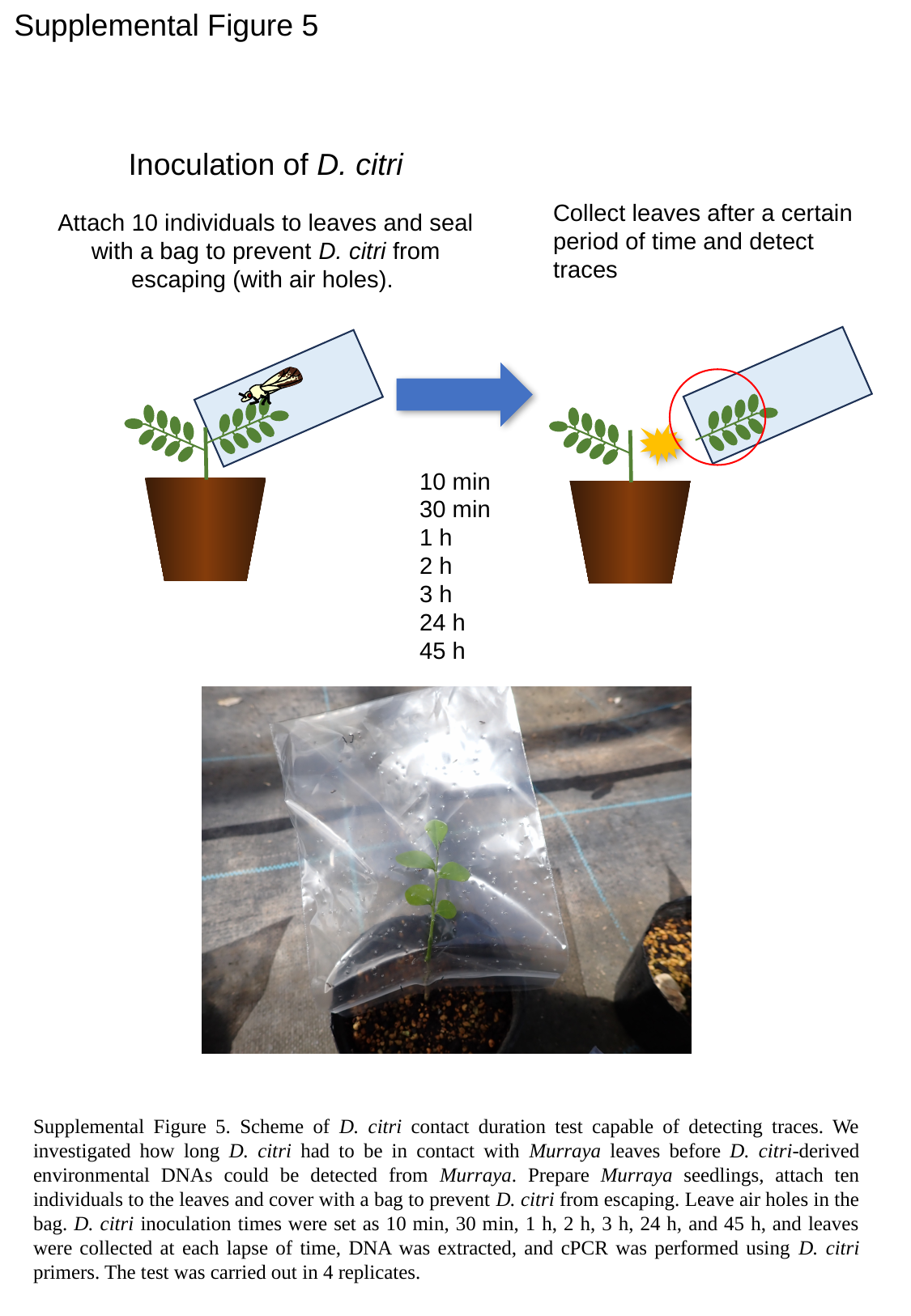

Supplemental Figure 5
Inoculation of D. citri
Collect leaves after a certain period of time and detect traces
Attach 10 individuals to leaves and seal with a bag to prevent D. citri from escaping (with air holes).
10 min
30 min
1 h
2 h
3 h
24 h
45 h
Supplemental Figure 5. Scheme of D. citri contact duration test capable of detecting traces. We investigated how long D. citri had to be in contact with Murraya leaves before D. citri-derived environmental DNAs could be detected from Murraya. Prepare Murraya seedlings, attach ten individuals to the leaves and cover with a bag to prevent D. citri from escaping. Leave air holes in the bag. D. citri inoculation times were set as 10 min, 30 min, 1 h, 2 h, 3 h, 24 h, and 45 h, and leaves were collected at each lapse of time, DNA was extracted, and cPCR was performed using D. citri primers. The test was carried out in 4 replicates.
