## Supplementary Figure 6 for "Investigation of host plant contact with *Diaphorina citri* (Hemipreta: Psyllidae) by detecting *D. citri*-derived environmental DNA": 23. Suppl. Figure 6 duration.pptx

### Slide 1
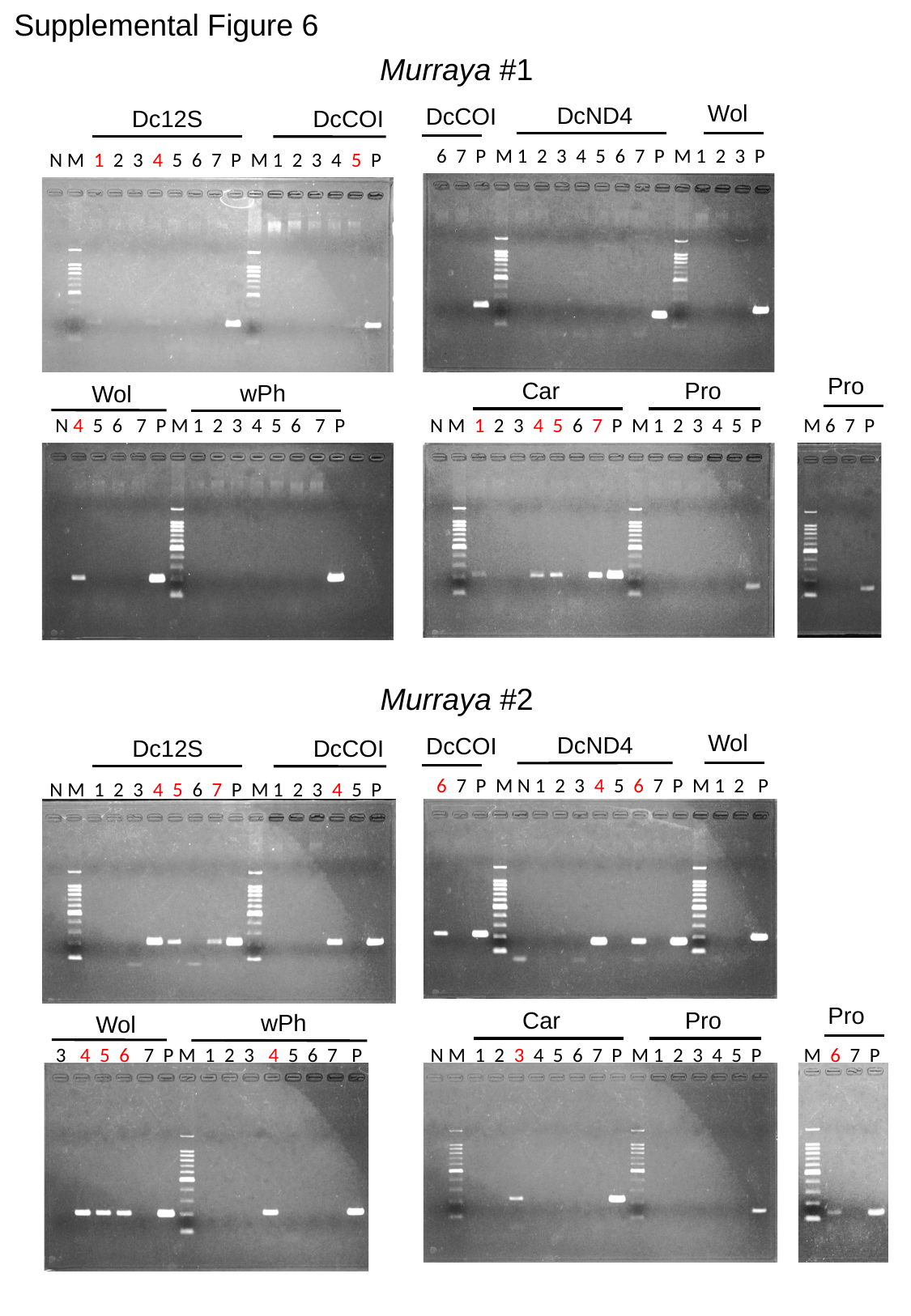

Supplemental Figure 6
Murraya #1
Wol
DcND4
DcCOI
DcCOI
Dc12S
6 7 P M 1 2 3 4 5 6 7 P M 1 2 3 P
N M 1 2 3 4 5 6 7 P M 1 2 3 4 5 P
Pro
Car
Pro
wPh
Wol
N 4 5 6 7 P M 1 2 3 4 5 6 7 P
N M 1 2 3 4 5 6 7 P M 1 2 3 4 5 P
M 6 7 P
Murraya #2
Wol
DcND4
DcCOI
DcCOI
Dc12S
6 7 P M N 1 2 3 4 5 6 7 P M 1 2 P
N M 1 2 3 4 5 6 7 P M 1 2 3 4 5 P
Pro
Car
Pro
wPh
Wol
3 4 5 6 7 P M 1 2 3 4 5 6 7 P
N M 1 2 3 4 5 6 7 P M 1 2 3 4 5 P
M 6 7 P

### Slide 2
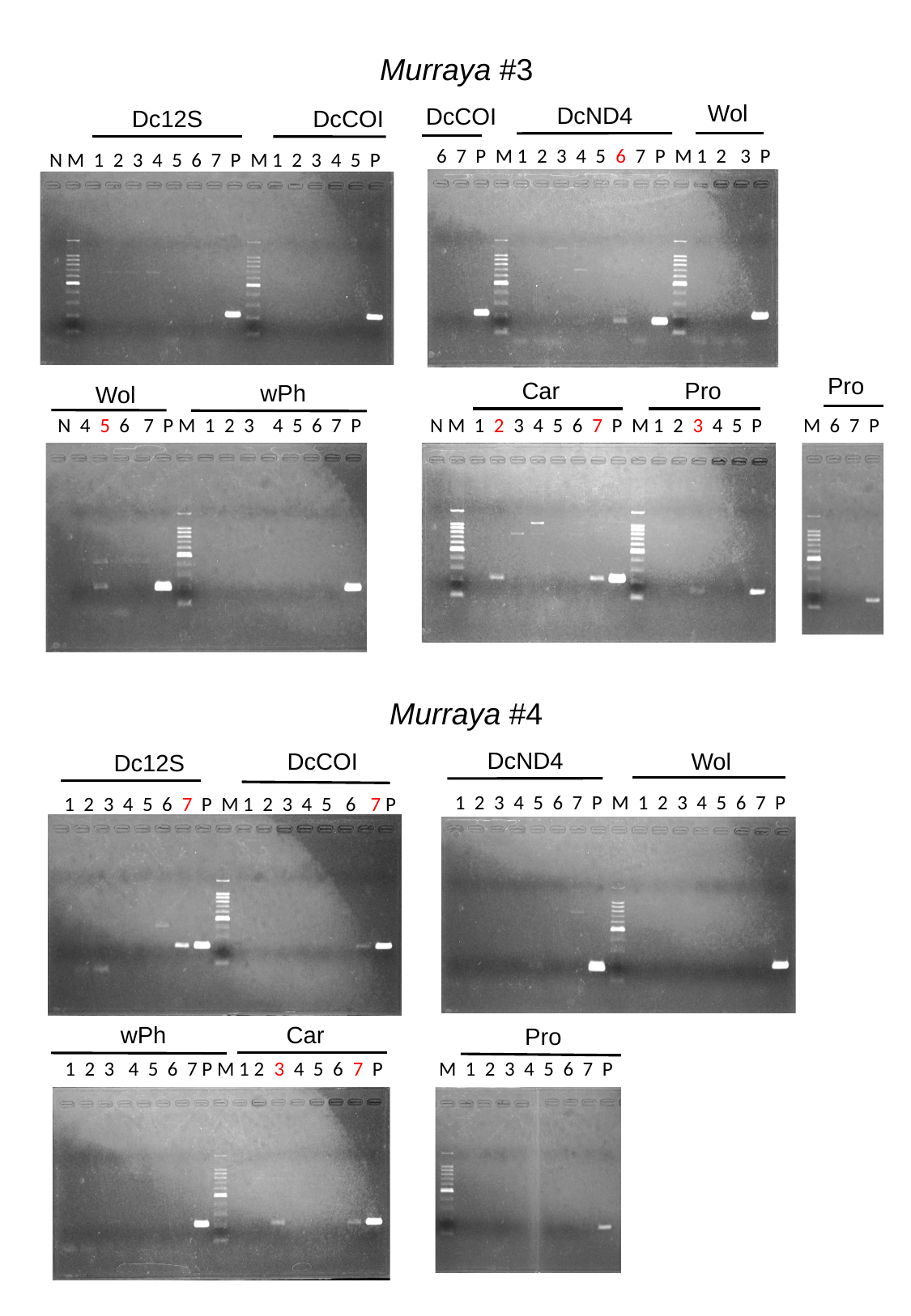

Murraya #3
Wol
DcND4
DcCOI
DcCOI
Dc12S
6 7 P M 1 2 3 4 5 6 7 P M 1 2 3 P
N M 1 2 3 4 5 6 7 P M 1 2 3 4 5 P
Pro
Car
Pro
wPh
Wol
N 4 5 6 7 P M 1 2 3 4 5 6 7 P
N M 1 2 3 4 5 6 7 P M 1 2 3 4 5 P
M 6 7 P
Murraya #4
DcND4
Wol
DcCOI
Dc12S
1 2 3 4 5 6 7 P M 1 2 3 4 5 6 7 P
1 2 3 4 5 6 7 P M 1 2 3 4 5 6 7 P
wPh
Car
Pro
1 2 3 4 5 6 7 P M 1 2 3 4 5 6 7 P
M 1 2 3 4 5 6 7 P

### Slide 3
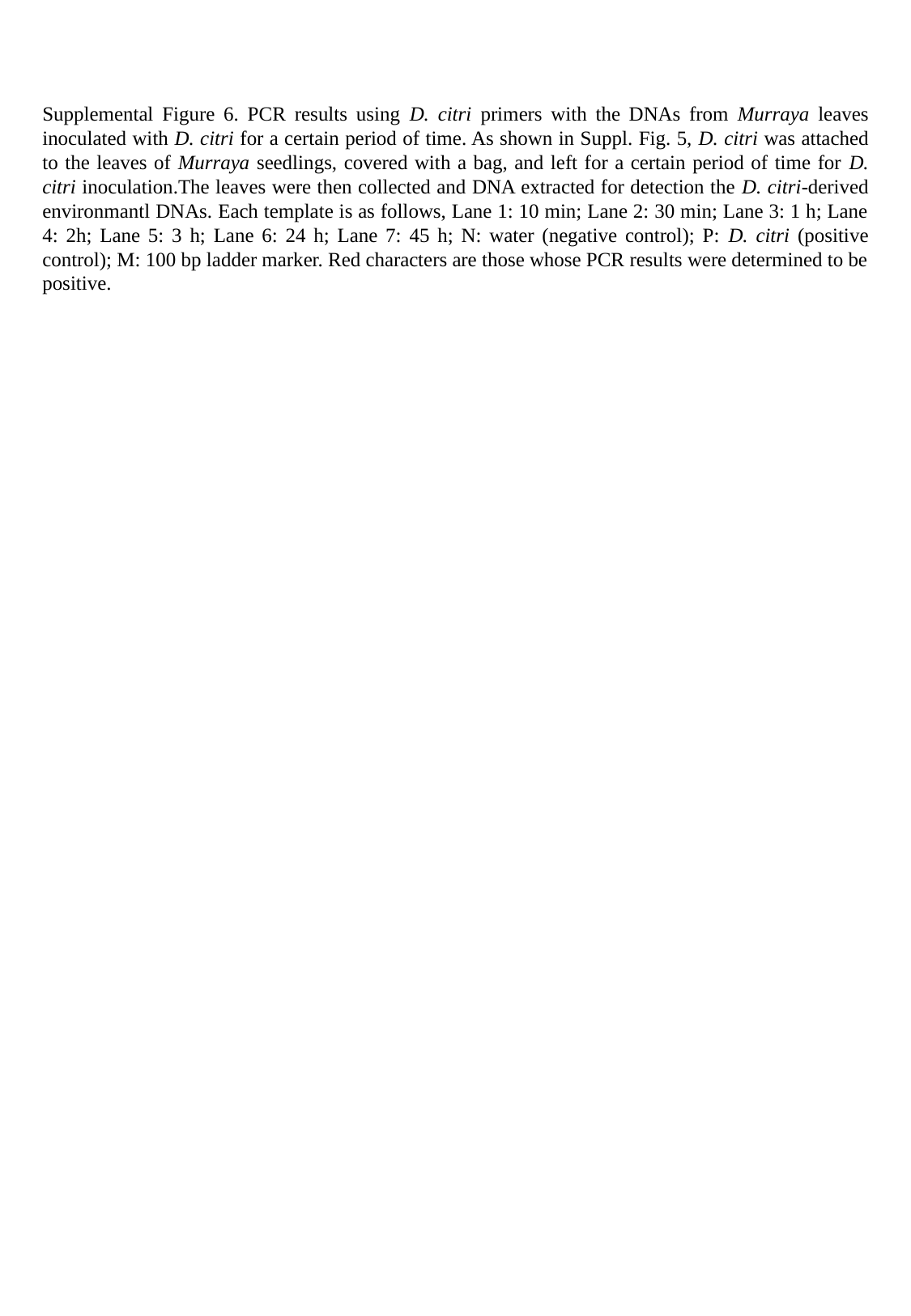

Supplemental Figure 6. PCR results using D. citri primers with the DNAs from Murraya leaves inoculated with D. citri for a certain period of time. As shown in Suppl. Fig. 5, D. citri was attached to the leaves of Murraya seedlings, covered with a bag, and left for a certain period of time for D. citri inoculation.The leaves were then collected and DNA extracted for detection the D. citri-derived environmantl DNAs. Each template is as follows, Lane 1: 10 min; Lane 2: 30 min; Lane 3: 1 h; Lane 4: 2h; Lane 5: 3 h; Lane 6: 24 h; Lane 7: 45 h; N: water (negative control); P: D. citri (positive control); M: 100 bp ladder marker. Red characters are those whose PCR results were determined to be positive.
