## Supplementary Figure 7 for "Investigation of host plant contact with *Diaphorina citri* (Hemipreta: Psyllidae) by detecting *D. citri*-derived environmental DNA": 24. Suppl. Figure 7 residual scheme-ed1.pptx

### Slide 1
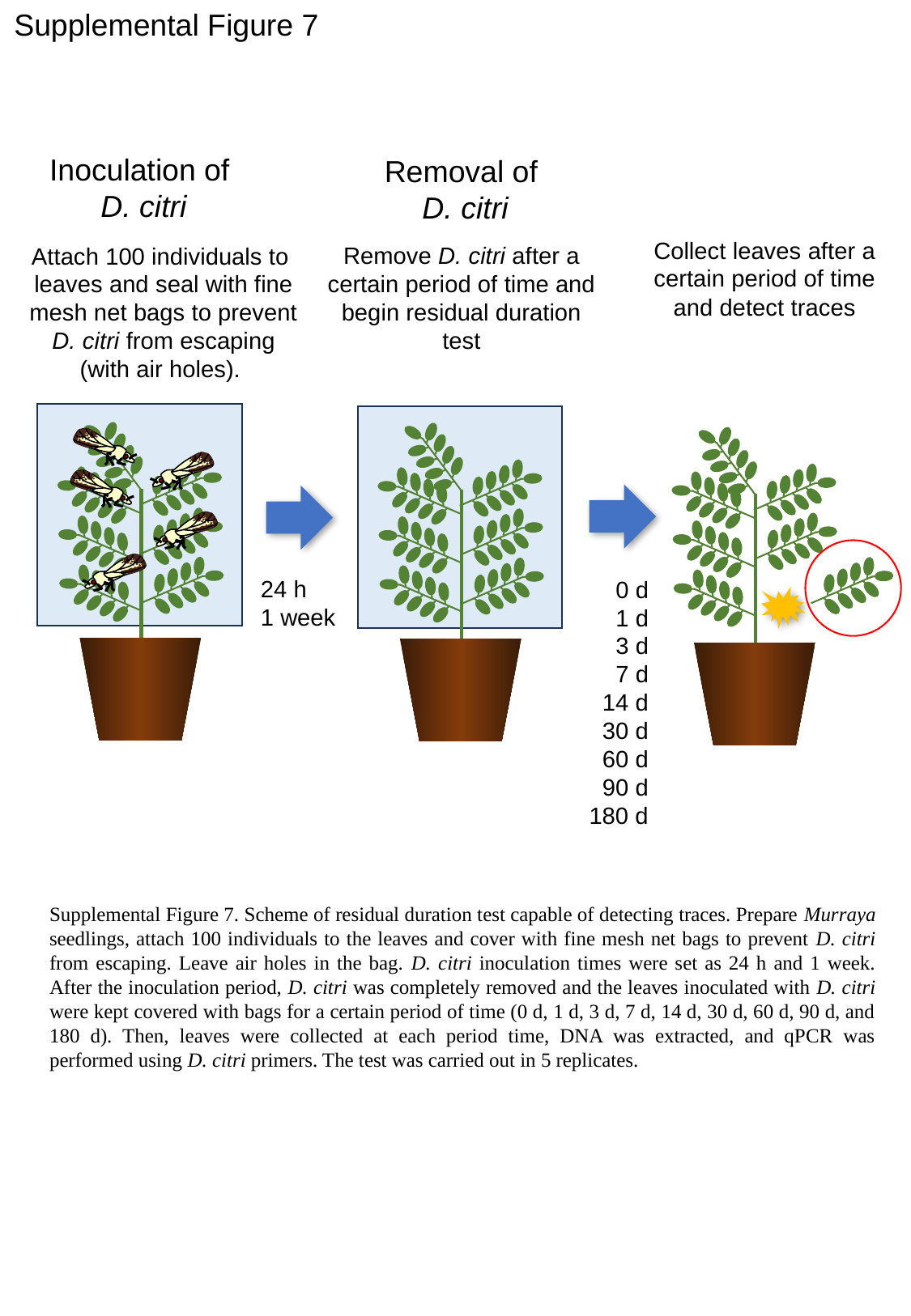

Supplemental Figure 7
Inoculation of
D. citri
Removal of
D. citri
Collect leaves after a certain period of time and detect traces
Remove D. citri after a certain period of time and begin residual duration test
Attach 100 individuals to leaves and seal with fine mesh net bags to prevent D. citri from escaping (with air holes).
24 h
1 week
 0 d
 1 d
 3 d
 7 d
 14 d
 30 d
 60 d
 90 d
180 d
Supplemental Figure 7. Scheme of residual duration test capable of detecting traces. Prepare Murraya seedlings, attach 100 individuals to the leaves and cover with fine mesh net bags to prevent D. citri from escaping. Leave air holes in the bag. D. citri inoculation times were set as 24 h and 1 week. After the inoculation period, D. citri was completely removed and the leaves inoculated with D. citri were kept covered with bags for a certain period of time (0 d, 1 d, 3 d, 7 d, 14 d, 30 d, 60 d, 90 d, and 180 d). Then, leaves were collected at each period time, DNA was extracted, and qPCR was performed using D. citri primers. The test was carried out in 5 replicates.
