## Supplementary Figure 8 for "Investigation of host plant contact with *Diaphorina citri* (Hemipreta: Psyllidae) by detecting *D. citri*-derived environmental DNA": 25. Suppl. Figure 8 cPCR Okinoerabu-ed1.pptx

### Slide 1
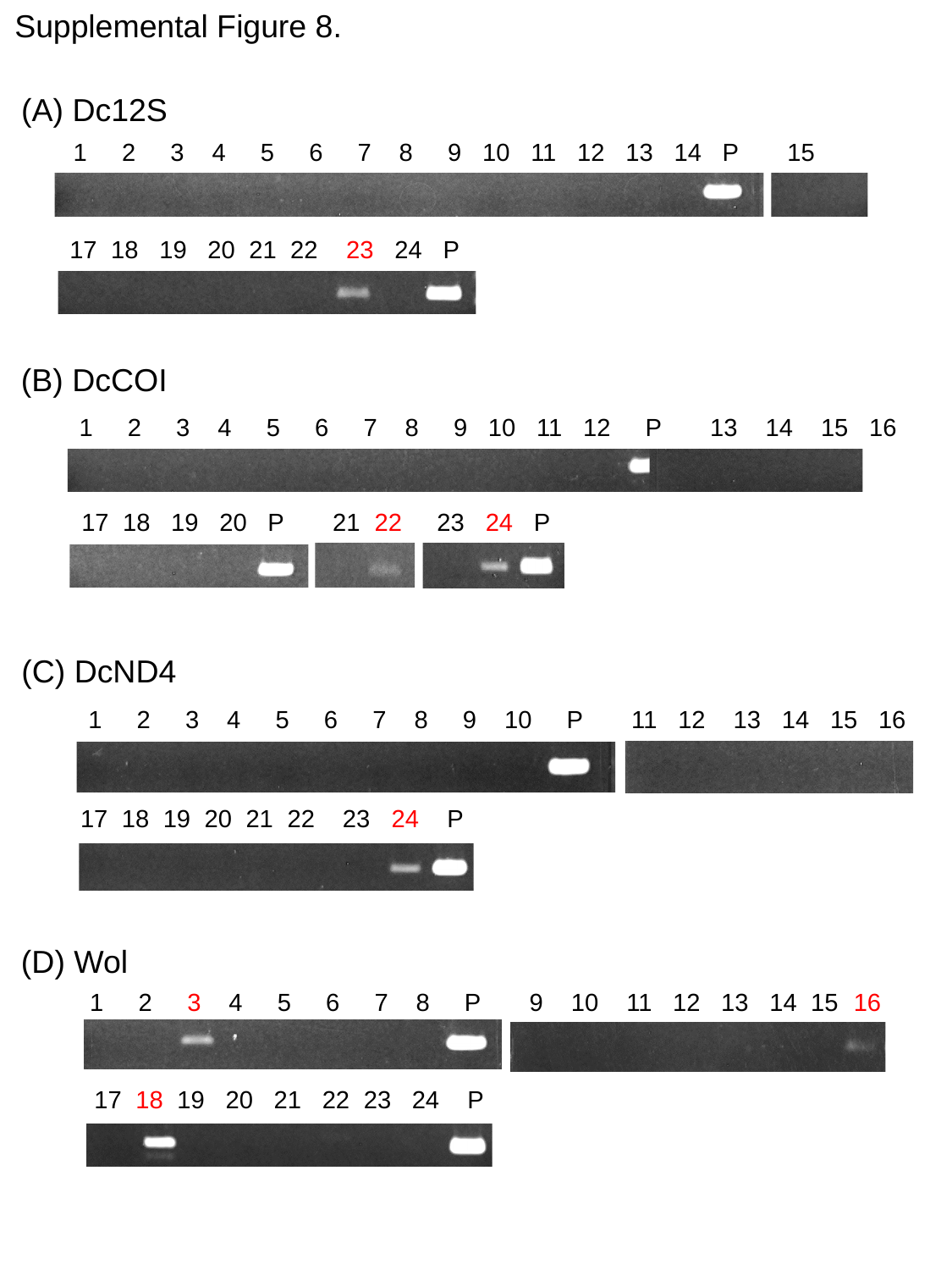

Supplemental Figure 8.
(A) Dc12S
1 2 3 4 5 6 7 8 9 10 11 12 13 14 P 15 16
17 18 19 20 21 22 23 24 P
(B) DcCOI
1 2 3 4 5 6 7 8 9 10 11 12 P 13 14 15 16
17 18 19 20 P 21 22 23 24 P
(C) DcND4
1 2 3 4 5 6 7 8 9 10 P 11 12 13 14 15 16
17 18 19 20 21 22 23 24 P
(D) Wol
1 2 3 4 5 6 7 8 P 9 10 11 12 13 14 15 16
 17 18 19 20 21 22 23 24 P

### Slide 2
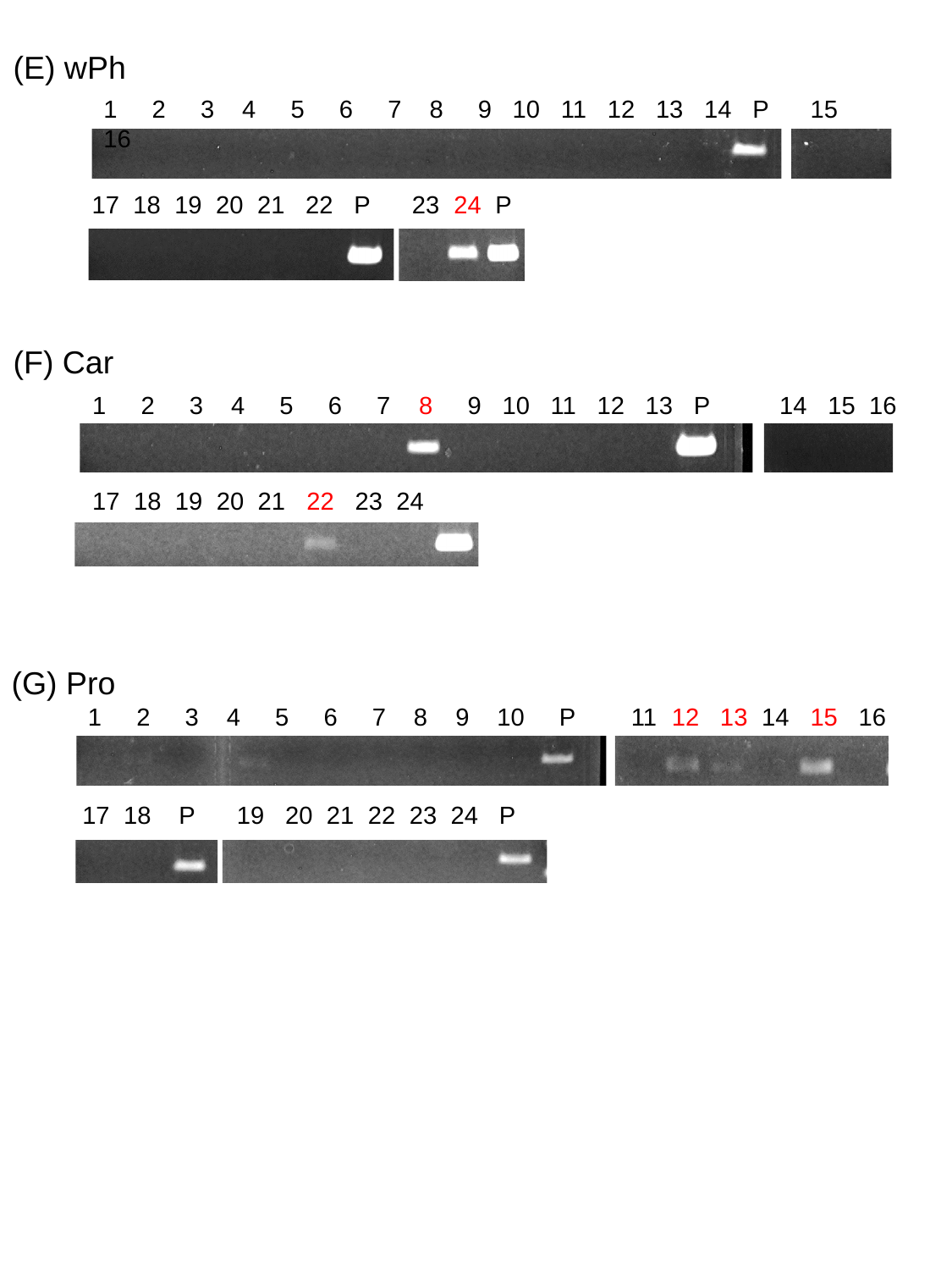

(E) wPh
1 2 3 4 5 6 7 8 9 10 11 12 13 14 P 15 16
17 18 19 20 21 22 P 23 24 P
(F) Car
1 2 3 4 5 6 7 8 9 10 11 12 13 P 14 15 16
17 18 19 20 21 22 23 24
(G) Pro
1 2 3 4 5 6 7 8 9 10 P 11 12 13 14 15 16
17 18 P 19 20 21 22 23 24 P

### Slide 3
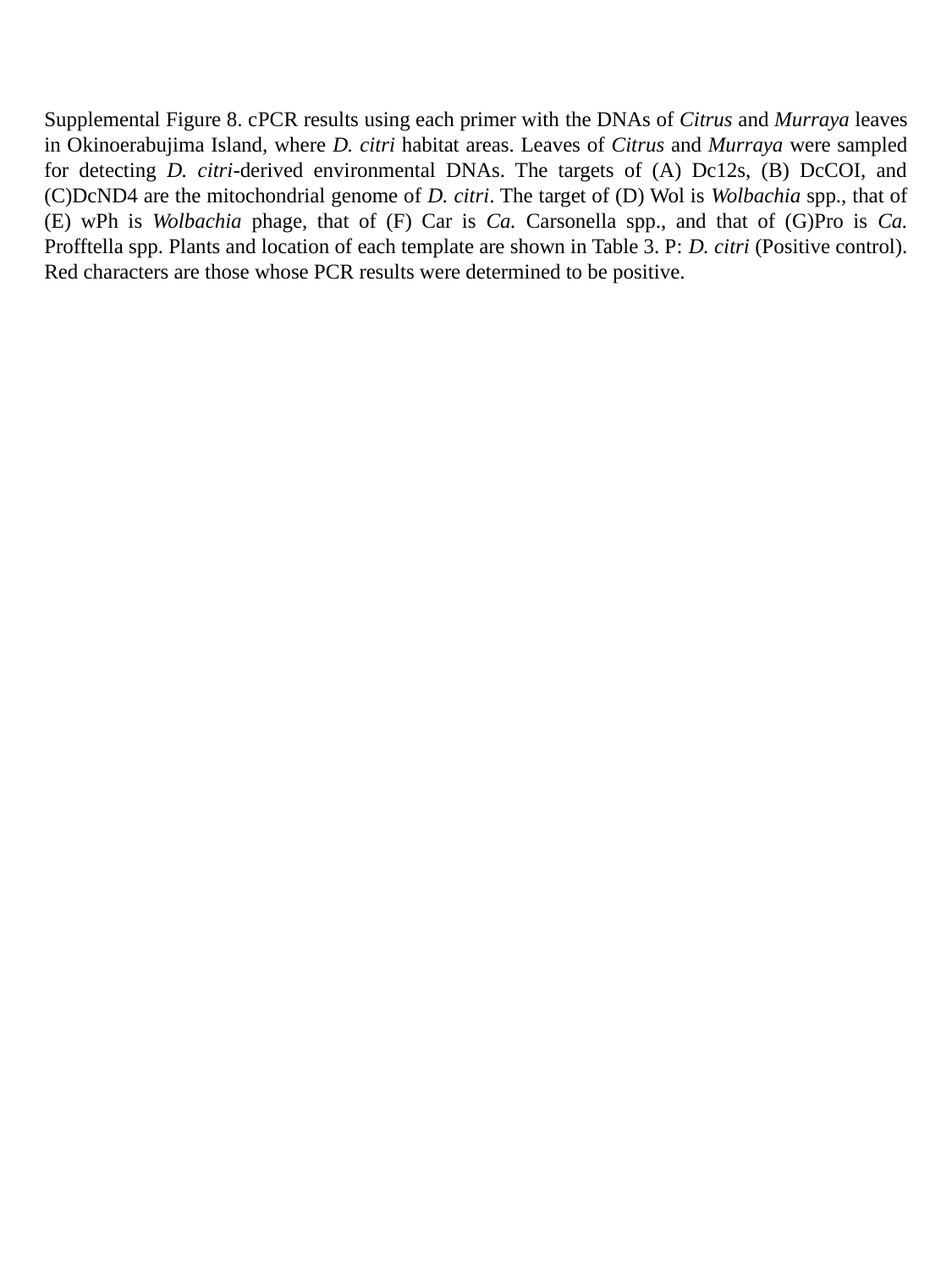

Supplemental Figure 8. cPCR results using each primer with the DNAs of Citrus and Murraya leaves in Okinoerabujima Island, where D. citri habitat areas. Leaves of Citrus and Murraya were sampled for detecting D. citri-derived environmental DNAs. The targets of (A) Dc12s, (B) DcCOI, and (C)DcND4 are the mitochondrial genome of D. citri. The target of (D) Wol is Wolbachia spp., that of (E) wPh is Wolbachia phage, that of (F) Car is Ca. Carsonella spp., and that of (G)Pro is Ca. Profftella spp. Plants and location of each template are shown in Table 3. P: D. citri (Positive control). Red characters are those whose PCR results were determined to be positive.
